# MINFLUX dissects the unimpeded walking of kinesin-1

**DOI:** 10.1101/2022.07.25.501426

**Authors:** Jan O. Wolff, Lukas Scheiderer, Tobias Engelhardt, Johann Engelhardt, Jessica Matthias, Stefan W. Hell

## Abstract

We report on an interferometric MINFLUX microscope that records protein movements with down to 1.7 nm precision within less than 1 ms. While such spatio-temporal resolution has so far required linking a strongly scattering 30-500 nm diameter bead to the much smaller protein, MINFLUX localization requires the detection of only down to 20 photons from an ~1-nm sized fluorophore. Harnessing this resolution, we dissect the unhindered stepping of the motor protein kinesin-1 on microtubules at up to physiological ATP concentrations. By attaching the fluorophore to different kinesin-1 sites and resolving steps and substeps of these protein constructs, we uncover a three-dimensional orientation change of the unbound kinesin head. We also find that kinesin-1 takes up ATP while only one head is bound, whereas hydrolysis of ATP occurs with both heads bound to the microtubule, resolving a long-standing conundrum of its mechanochemical cycle. Our results establish MINFLUX as a non-invasive tool for tracking protein movements and probing submillisecond structural rearrangements with nanometer resolution.

---

Exploring movements and conformational changes of proteins lies at the heart of unraveling the inner workings of a cell, but the tools for accomplishing this task have so far been limited. Nanometer-sized protein motions of millisecond duration can be retrieved by tethering the protein to a bead held in an infrared optical trap, and measuring the bead movement in the trap (*1–3*). However, this preparation subjects the protein to a force and hence does not allow the direct observation of its entirely free motion. Besides, the 80-500 nm diameter beads required for optical trapping are by orders of magnitude larger than the protein itself, which entails its own limits, including susceptibility to laser-induced heating (*4*). Alternatively, a protein can be observed with no or minimal restrictions by labeling it with an ~1-nm-sized organic fluorophore and recording its motion with the camera of a light microscope (*5,6*). The position of the label is then inferred from the peak of the fluorescence diffraction pattern rendered by *N* camera-detected photons per position. Unfortunately, the resulting localization precision *σ* scales with 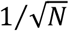, meaning that *σ*=1-2 nm typically requires *N* > 2500 photons (*7*). Thus, even the brightest fluorophores entail localization times of the order of hundred milliseconds. Camera-based localization therefore cannot live up to the spatio-temporal resolution (STR) provided by optical traps. Replacing the tiny fluorophore with a laser scattering gold bead of >30 nm diameter (*8–10*) can compensate for this shortfall, but then again the gold bead volume, drag and its electrostatic interactions bring back the question on whether the protein motion is unimpeded.

Not surprisingly, these limits are also reflected in the understanding of the arguably best studied moving protein, the homodimeric motor protein kinesin-1, hereafter called kinesin, which is responsible for the anterograde transport on microtubules, and whose malfunction is linked to a number of diseases (*11–14*). Although all the above tools have substantially advanced our understanding of how kinesin walks, many details of its mechanochemical cycle have remained controversial or elusive (*15, 16*).

We reasoned that MINFLUX (*17*), a recently introduced microscopy method for localizing fluorophores with a minimal - rather than maximal - number of detected photons *N*, should greatly improve the study of protein movements. For a given *N*, MINFLUX (*17–20*) typically renders a STR of 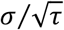 with about 10-fold improved *σ*, or a 100-fold increased temporal resolution *τ* compared to popular camera-based localizations (*18*). Thus, using a single fluorophore of about 1 nm in size, a STR is attained that has so far required the use of bulky beads. This unique combination of STR and a tiny label has motivated us to revisit the walking of kinesin. Here, we report on an interferometric MINFLUX implementation that delivers submillisecond/nanometer STR in protein tracking. Harnessing this STR, we dissected the steps and substeps of the heads and the coiled-coil domain of kinesin. The direct observation of unhindered substeps allowed us to derive in which state adenosine-5’-triphosphate (ATP) binds and hydrolyzes, as well as to uncover orientation changes of functional subunits of kinesin during stepping. Our study concomitantly establishes MINFLUX as a tool for examining fast protein movements at the very nanometer scale with minimal or no impediment.

## Interferometric MINFLUX maximizes fluorophore localization precision

A scanning MINFLUX microscope features a beam for fluorophore excitation having a central intensity minimum (‘zero’) that is positioned in the sample with subnanometer-precision and a confocal point detector for measuring the emitted photons (Fig. 1A). The closer the central excitation minimum is to the fluorophore, the lower is the fluorescence rate, meaning that the number of detections readily discloses the distance between the unknown position of the fluorophore and the well-known position of the minimum. In fact, the intensity of the MINFLUX excitation beam around the minimum increases, in good approximation, quadratically with distance to the minimum (Fig. 1B), with steepness depending on the beam’s focusing angle, wavelength and power. Consequently, the fluorescence detection rate (per unit beam power) displays the same quadratic dependence on the fluorophore-to-minimum distance (Fig. 1B). If the rate is minimal, i.e. down to background level, the fluorophore is localized, because the positions of the fluorophore and that of the excitation minimum are identical. However, due to the derogatory role of background, matching the two positions at the Ångström level is usually not possible. Fortunately, such perfect matching is not needed, because the matching point and hence the fluorophore position can be precisely derived from a small number *N* of photons gained by targeting the minimum to two or more positions within a spatial interval *L* containing the fluorophore (Fig. 1B).

**Fig. 1.**
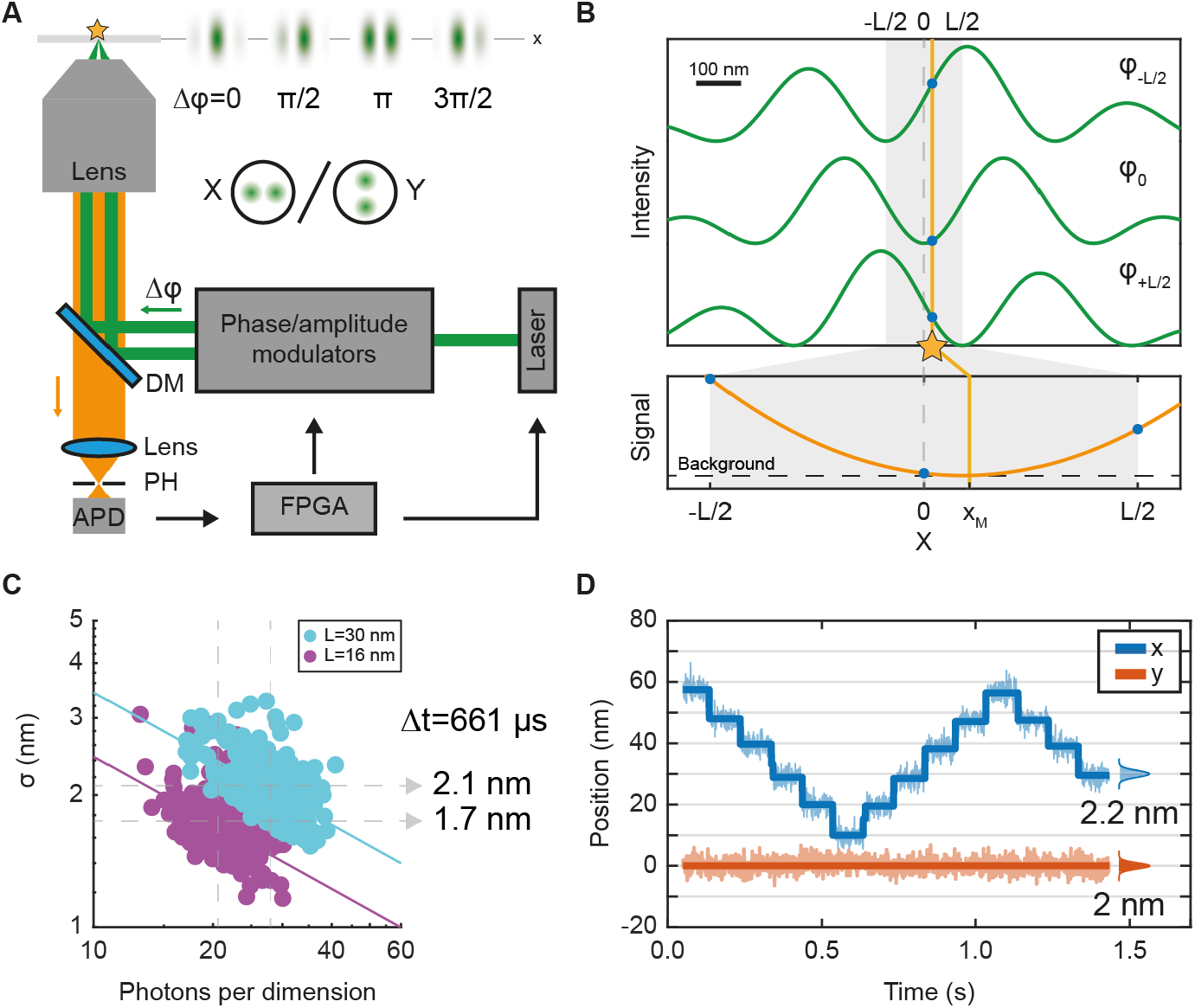
Interferometric MINFLUX microscope provides nanometer localization precision of a fluorophore with 20-40 detected photons. **(A)** Simplified setup: a 640 nm laser beam is shaped by a phase modulator and an amplitude modulator to create a pair of beamlets with a defined phase difference ϕ in the entrance pupil of an objective lens. Their interference creates an intensity pattern with a local line-shaped minimum in the focal plane. The minimum is shifted by changing ϕ with the phase/amplitude modulator. Two orthogonal pairs of beams are used for covering both the x- and the y-direction. The fluorescence collected by the lens passes a dichroic mirror (DM), that otherwise deflects the laser light, is focused onto a confocal pinhole (PH) and detected by an avalanche photodiode (APD). The (x,y) localization algorithm is implemented in a field programmable gate array (FPGA) directing the minimum to specified (x,y) positions, depending on the number of photons detected by the APD. **(B)** Top: MINFLUX localization in one dimension, using a change of ϕ to place the minimum at positions –L/2, 0 and L/2 of a linear interval around the expected molecule position *x*_*M*_. Bottom: The photons counts measured with the minimum at these three points allow retrieving *x*_*M*_ by fitting with a parabola. In iterative MINFLUX localizations where the excitation intensity minimum approaches the fluorophore, the laser power is increased accordingly to keep the fluorescence rate at the same level. Background is due to non-vanishing excitation intensity at the minimum, stray light and detector dark counts. **(C)** Localization precision *σ* of single surface-immobilized ATTO647N fluorophores using L=30 nm (263 fluorophores) and L=16 nm (232 fluorophores), yielding *σ*= 2.1 nm and 1.7 nm, respectively. **(D)** Tracking of a single fluorophore moved by a piezo-electric stage (L=30 nm, 2 nm residual noise, 0.607 ms temporal resolution, 70 photons per (x,y)-localization).

Therefore, typical MINFLUX localization is performed iteratively (*19*) by continually shifting the minimum closer to the fluorophore. Normally, the localization starts out from an interval *L* of about the diffraction limit (200 nm), which is then reduced on the basis of the initially derived precision *σ*_0_. In theory, an iterative reduction of *L* in proportion to the precision *σ*_*k*−1_ of the previous step, *L*_*k*_ = *ασ*_*k*−1_, gives rise to an exponential increase in precision after *k* steps: 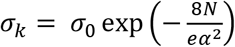. The regularization parameter *α* ensures that the next *L*_*k*_ is small enough to quickly zoom-in on the fluorophore, but large enough to keep the fluorophore in the interval. This exponential increase in precision with *N* signifies a most efficient use of the detected photons and should be contrasted with the sluggish 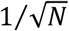 dependence in camera-based localization (see supplementary text 2.2). The iterative reduction of *L*_*k*_ ends just before the quadratic dependence disappears amid background. Hence, practical MINFLUX localization precision *σ* is limited by the steepness-to-background ratio of the excitation beam. In principle, steepness can be arbitrarily increased by increasing the beam power, but since this measure also increases the background, we designed a MINFLUX system that inherently yields higher steepness compared to reported donut-based systems (*17–20*).

Specifically, our MINFLUX setup featured two pairs of oblique beams that interfered destructively in the focal plane (Fig. 1A). While one of the pairs was arranged in the x-direction, rendering a y-oriented line-shaped (LS) minimum for x-localization, the pair for y-localization was arranged accordingly in y-direction. LS minima have also been used in STED microscopy (*21*), because they require fewer polarization and aberration optimizations, while providing higher steepness (fig. S1) and lower background. Altering the phase difference of the x-arranged beams, moved the y-oriented LS minimum with Ångström precision in the x-direction, and vice versa. By targeting the minima to coordinates by –*L*_*k*_/2, 0 and *L*_*k*_/2 away from the last estimated fluorophore position, the position was iteratively established for each dimension (x and y), based on the number of detections (Fig. 1B). The (x,y) trajectory was obtained by repeatedly switching between x and y using an electro-optical modulator (fig. S2 and fig. S3). Once *L*_*k*_ =16 nm was reached, as few as ~20 detected photons sufficed to localize single immobilized ATTO 647N fluorophores with an average precision *σ*=1.7 nm per dimension (Fig. 1C). For *L*_*k*_ =30 nm, a precision *σ*=2.1 nm was obtained with ~28 photons. Since the average signal-to-background ratio (SBR) was more than 3 times higher for *L*_*k*_ =30 nm, we carried out all tracking measurements with *L*_*k*_ *=L*=30 nm (fig. S4), ensuring higher robustness in the process. Note that in fluorophore tracking, the successive small changes in fluorophore position inherently allowed for the continual use of *L* =30 nm and hence for the maximal use of the *N* photons detected.

The tracking accuracy of our MINFLUX system is highlighted by moving an individual ATTO 647N fluorophore on a periodic stepping trajectory along the x-axis of a piezo-electric stage (Fig. 1D). The steps were fitted with an algorithm based on an iterative change-point search (*22*), which was used throughout our study. Our analysis showed that ~70 photons recorded within 607 μs clearly identified the steps with σ=2 nm in both x- and y-direction.

## MINFLUX observes substeps and coiled-coil rotation of kinesin

Under consumption of an ATP molecule, the catalytic motor domains (heads) of kinesin take hand-over-hand steps of 16 nm (regular steps) amounting to twice the tubulin dimer spacing. Their conjoining coiled-coil domain is thus translocated in discrete 8 nm steps (*1, 3, 6, 23*). However, whether this mechanism is symmetric or asymmetric is still debated (*24–26*). Camera-localization-based Fluorescence Imaging with One-Nanometer Accuracy (FIONA) (*5*) resolved regular kinesin steps using a single fluorophore label at one of the heads, but its time resolution of several hundreds of milliseconds required slowing down movement through administering ATP concentrations that were about thousand times lower than in a cell (*6*). As a matter of fact, addressing substeps at physiological ATP concentrations has so far required the use of beads.

For example, an optical trap study recently observed force-dependent substeps by tracking a germanium bead of ~80 nm diameter attached to the kinesin coiled-coil domain (*27*). Thus, like in all optical trap experiments, only the protein center-of-mass movements and not those of individual heads could be examined. Attaching a gold bead to a kinesin head allowed tracking these movements. However, different studies came to opposing results regarding the long-standing question of when ATP is bound (*15, 16*). In fact, simulations (*28, 29*) suggest that this discrepancy is due to the different labeling positions, as the beads are >200 times larger in volume than the kinesin head.

Harnessing MINFLUX, we first investigated the stepping of different cysteine-light truncated kinesin constructs labeled with a fluorophore at various protein positions via maleimide coupling. The kinesin molecules were introduced into a flow cell in which biotinylated and fluorescently labeled (Alexa Fluor 488) microtubules were attached via neutravidin to a PLL-PEG-biotin polymer coated coverslip. For kinesin center-of-mass tracking, we labeled construct N356C at its solvent-exposed cysteine introduced in the coiled-coil domain (Fig. 2A).

**Fig. 2.**
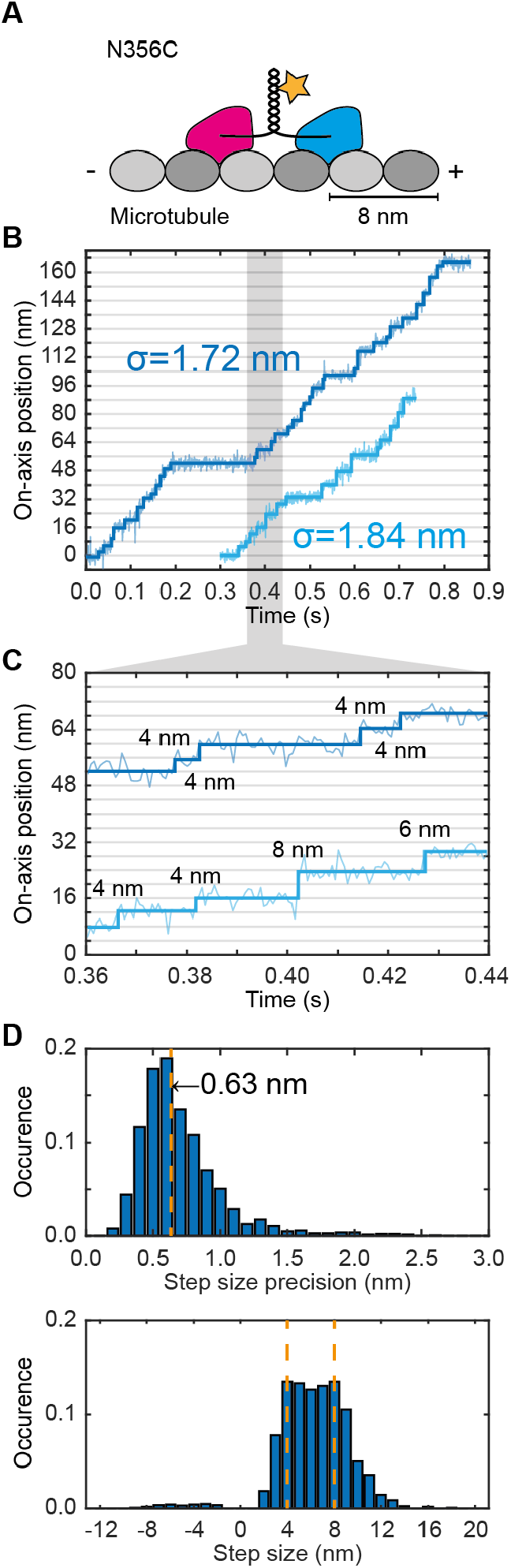
MINFLUX tracking of kinesin exhibits 4 nm center-of-mass substeps. **(A)** Scheme of kinesin walking on a microtubule indicating the labeling position of the fluorophore in the coiled-coil domain. **(B)** Exemplary position traces recorded along the microtubule axis at 10 μM ATP. The data are overlaid with the detected step function shown as thick darker lines. **(C)** Zoom-in of the traces shown in (B) between 0.36 s and 0.44 s as highlighted by grey shading. **(D)** Histograms of the step size precision (top) and the step sizes (bottom) for N=1739 steps. The median step size precision is 0.63 nm. Orange dashed lines highlight 4-nm-sized substeps and 8-nm-sized regular steps.

We recorded 1D-traces of individual kinesin dimers labeled with a single fluorophore (degree of labeling of 1, DOL1) walking along the microtubule axis (on-axis displacement) with a temporal resolution of ~1 ms and a residual position noise of *σ*≈1.8 nm (Fig. 2B and C). These initial measurements were performed at 10 μM ATP concentration entailing a walking speed of ~280 nm/s. The traces were recorded with run lengths up to ~180 nm. Based on the residual noise and the number of localizations between steps, we determined a median precision of the measured step size of 0.63 nm (Fig. 2D). A histogram of all kinesin center-of-mass steps revealed a size range of about 3-11 nm (Fig. 2D), with equally high peaks at 8 nm and 4 nm, corresponding to expected regular steps and substeps, respectively. Note that the latter have so far not been observed without attaching a much bigger bead to the protein (*27*).

With the same kinesin construct, we also recorded traces at physiological 1 mM ATP concentration. Despite the now increased walking speed of about 550 nm/s, both regular steps and substeps of the kinesin center-of-mass were resolved (fig. S5). The substantially smaller fraction of detected substeps indicated a reduced detection efficiency due to shorter substep dwell times (fig. S6). Strikingly, the step size histogram did not exhibit its maximum at 8 nm as expected for regular steps, but showed an unexpectedly high occurrence of 6 nm and 10 nm steps (Fig. 3A top, DOL1). Plotting the sequence of consecutive step sizes in a 2D histogram revealed their sum frequently matching 16 nm, indicating that these unusual steps typically occurred sequentially (Fig. 3B top, DOL1). Since the non-zero radius of the coiled-coil domain (~1.0 nm, inferred from PDB 1D7M) and the distance between maleimide and fluorophore core (~1.0 nm, fig. S7) added up to a total fluorophore displacement of ~2 nm from the coiled-coil axis of kinesin and assuming the fluorophore displacement vector had a component parallel to the walking direction, we reasoned that the observed stepping asymmetry is due to a rotation of the coiled-coil domain during a regular step (Fig. 3C left, DOL1).

**Fig. 3.**
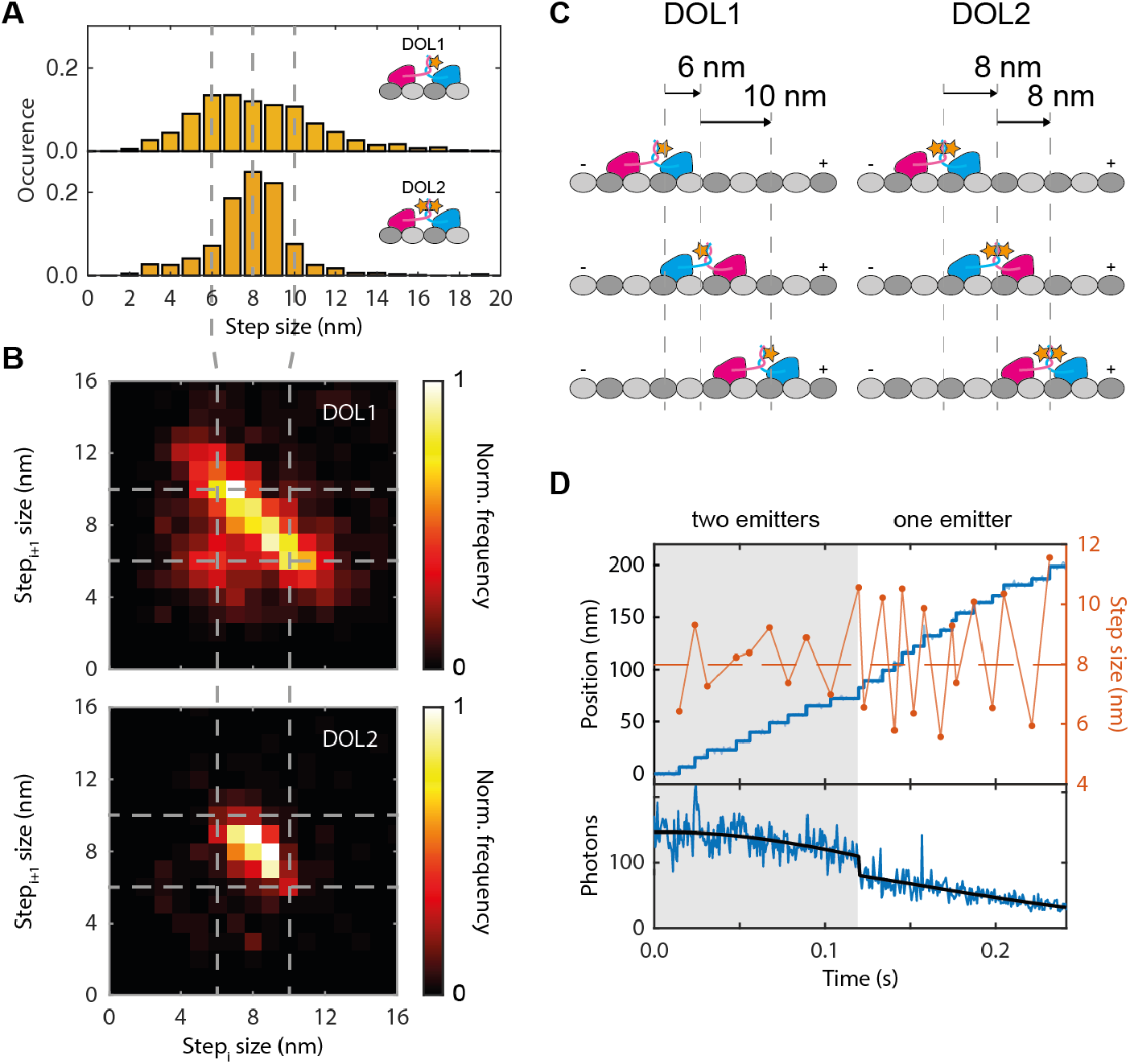
Rotation of the coiled-coil domain during kinesin stepping. **(A)** Step size histograms for 1D kinesin center-of-mass tracking with a single fluorophore (DOL1, N_DOL1_=1759) and two fluorophores on same protein site of the kinesin dimer (DOL2, N_DOL2_=625) at 1 mM ATP concentration. Relevant step sizes of 6 nm, 8 nm and 10 nm are highlighted by dashed grey lines. **(B)** 2D histogram of consecutive step sizes showing alternating 6 nm and 10 nm steps for DOL1 (top) and predominantly successive 8 nm steps for DOL2 (bottom). **(C)** Suggested model of coiled-coil domain rotation explaining the alternating sequence of larger and smaller steps at DOL1 and the true 8 nm coiled-coil displacement at DOL2. **(D)** Exemplary position trace of DOL2 (blue, top) with one emitter bleaching after ~0.12 s as evident from photon trace (blue, bottom) and Gaussian fit accounting for a bleaching step (black). Before the bleaching event, the step size (red) varies around 8 nm, whereas after bleaching the kinesin walking exhibits an asymmetric sequence of 10 nm and 6 nm steps. The overall Gaussian-shaped decrease in emission is caused by kinesin walking out of the stationary confocal volume.

To test this hypothesis, we labeled the same construct with an excess of fluorophores, ensuring that the cysteines at amino acid position 356 of both monomers each carried a fluorophore (degree of labeling of 2, DOL2) (fig. S7 and fig. S8). As a result, we found stepping symmetry reinstated, since MINFLUX inherently localized the midpoint between two adjacent identical fluorophores; note that in our sample design, this midpoint coincided with the coiled-coil axis (Fig. 3C right, DOL2). To ensure that the histogram of the DOL2 experiment exclusively represents steps of kinesins with both fluorophores emitting, only DOL2 tracking data (characterized by a photon rate > 125 kHz, as determined from the DOL1 data) were plotted. Supporting our hypothesis of a coiled-coil rotation, the resulting step size histogram indeed shows a rather narrow peak at 8 nm (Fig. 3A bottom, DOL2) and the 2D histogram of consecutive step sizes indicates the dominance of successive 8 nm steps (Fig. 3B bottom, DOL2). In a trace (Fig. 3D) where one of the fluorophores bleached (at ~0.12 s), a clear difference in the step sizes before and after bleaching becomes apparent: ~8 nm (before) and alternating ~10 nm and ~6 nm (after). We conclude that the coiled-coil domain significantly rotates when kinesin steps. Whether consecutive steps cause a unidirectional (*26, 30*) or a back-and-forth rotation (*24, 25*) cannot be deduced from this experiment alone.

## ATP binds in one-head-bound state

Next we explored whether ATP binds to kinesin in its one-head-bound state (1HB, only leading head bound) or its two-head-bound state (2HB, leading and trailing head associated with their binding site), a longstanding open question concerning the kinesin mechanochemical cycle (*15, 16, 31–34*). We employed the construct T324C labeled at its solvent-exposed cysteine (DOL1), which is located at the C-terminal end of the α6-helix, adjacent to the neck linker on the head (Fig. 4A). When the head is microtubule-bound, the label is in the center on the right side of the motor domain with respect to the walking direction. We recorded 2D-traces (on-axis and off-axis displacement) at ATP concentrations of 10 μM, 100 μM and 1 mM. By tracking one of the heads rather than the kinesin center-of-mass, we observed traces with regular steps of 16 nm, the distance between every second binding site on the microtubule, and substeps of 8 nm resulting from the labeled head occupying an unbound intermediate state (Fig. 4B). Accordingly, the on-axis step size histogram shows a fraction of regular steps peaking at 16 nm and a fraction of substeps distributed around 8 nm. In good agreement with the results obtained from construct N356C, the fraction of detected substeps decreased with increasing ATP concentration, indicating an ATP-dependence of the unbound state (Fig. 4C). Note the unexpectedly broad distribution of substep sizes, which is discussed further down.

**Fig. 4.**
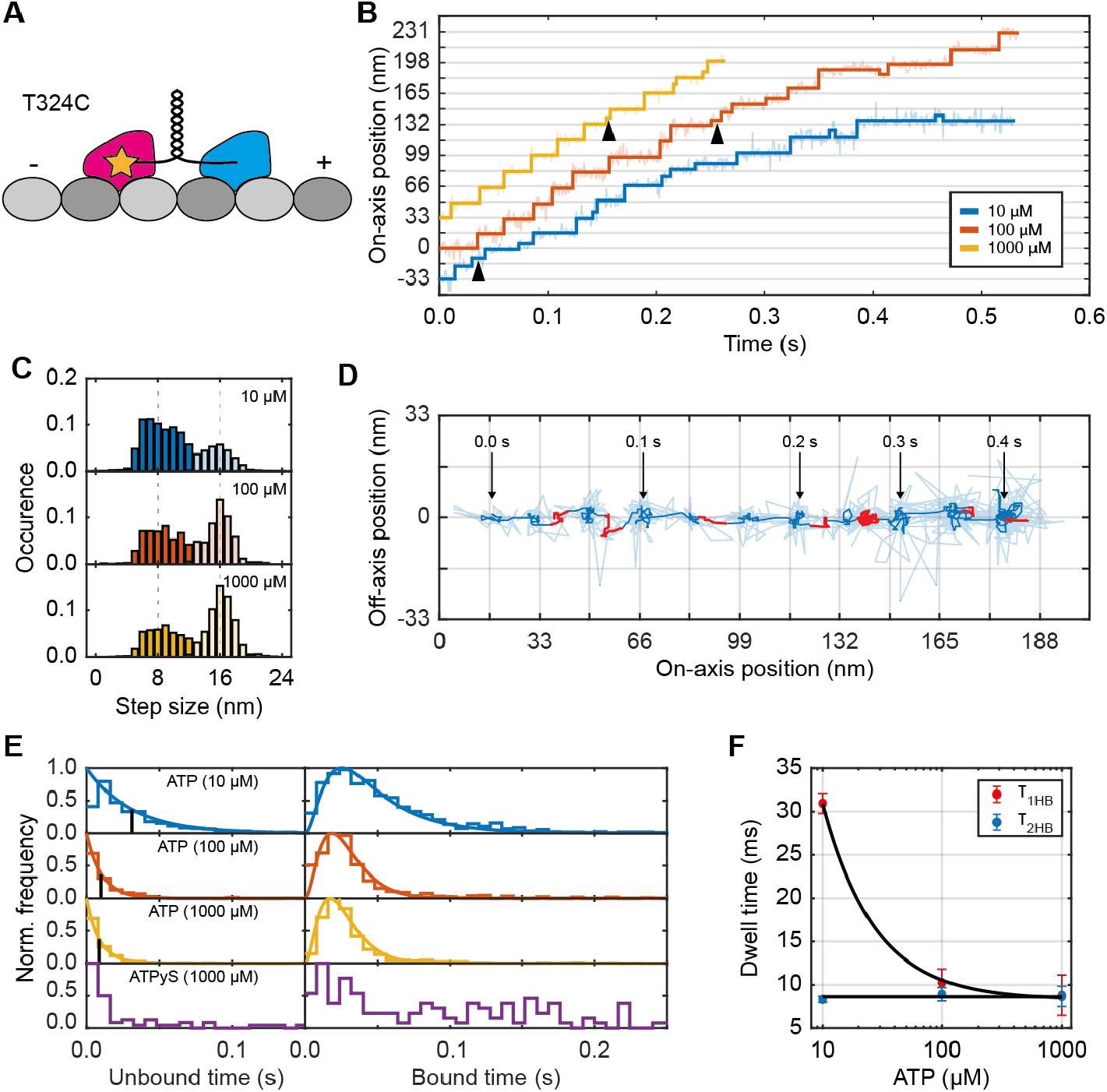
Kinesin awaits ATP binding in one-head-bound state. **(A)** Construct T324C with labeling position of the fluorophore on a kinesin head. **(B)** Exemplary traces recorded at 10 μM (blue), 100 μM (orange) and 1 mM (yellow) ATP concentration with distinct plateaus spaced by ~16 nm. Exemplary substep-plateaus between two 8 nm steps are highlighted by black arrows. **(C)** Histogram of detected step sizes for each ATP concentration showing 8 nm substeps (darker filling; N_10μM_=1160, N_100μM_=568, N_1mM_=892) and 16 nm regular steps (lighter filling; N_10μM_=558, N_100μM_=653, N_1mM_=1299) as assigned by the Hidden-Markov-Model (HMM). **(D)** 2D representation of the 10 μM trace shown in (B) with time stamps. Plateaus identified by the HMM as unbound states are highlighted in red. For improved visibility, the raw data points are overlaid with a 5 ms moving median filter. **(E)** Normalized histogram of residence times in the bound and unbound state as assigned by the HMM for different ATP concentrations (stair functions) together with the fitted model (colored lines) assuming a single time constant τ_1HB_ for the unbound state, and the sum of twice τ_2HB_ and once τ_1HB_ for the bound state. Black vertical lines indicate τ_1HB_ as resulting from the fit. **(F)** Comparison of τ_1HB_ and τ_2HB_ for different ATP concentrations. Black lines show the fitted Michaelis-Menten kinetics (K_M=_27±3 μM, k_ATP_=4.3±0.8 s^−1^μM^−1^) for the 1HB state and a constant fit (τ_2HB_=8.6 ms) for the 2HB state. Error bars denote the 64 % confidence interval of the fit.

To explicitly identify the bound states (B), in which the labeled kinesin head is located at its microtubule binding site, and unbound states (U), in which it is located in-between, we assumed five different state transitions and applied a Hidden Markov Model (HMM) to the sequence of detected step sizes (see Materials and Methods section 1.4.7). Most 8-nm steps correspond to transitions between the two states (B→U, U→B), but can also be caused by so-called slip states (B→B, see next section) in rare cases. Transitions between bound states (B→B), during which the intermediate unbound state was too short and hence missed, were assumed to be the major source of 16-nm steps. Potential 16-nm transitions between unbound states (U→U) of the labeled head comprise a series of the transitions explained above (labeled head U→B, unlabeled head B→U and U→B, labeled head B→U), so missing these states was assumed to be highly unlikely (see supplementary text 2.3).

Based on these premises, we assigned bound and unbound states to all traces of the kinesin construct T324C with the HMM (see representative trace in Fig. 4D). Using this assignment, the dwell times in the bound and unbound states were determined for each ATP concentration. To obtain the average dwell times *τ*_1*HB*_ and *τ*_2*HB*_ of the underlying 1HB and 2HB states, respectively, the histograms of residence time in the bound and unbound states were fitted simultaneously (Fig. 4E). The unbound state (1HB with labeled head unbound) data were fitted with a mono-exponential decay function. For the bound state (2HB with labeled head leading, 1HB with labeled head bound, 2HB with labeled head trailing), a combination of three exponential decay functions was used under the assumption of equal binding kinetics for both heads. Matching the trend deduced from the step size histograms in Fig. 4C, *τ*_1*HB*_ substantially increased with decreasing ATP concentrations from around 8 ms for 1 mM ATP to over 30 ms for 10 μM ATP (Fig. 4F).

In contrast, *τ*_2*HB*_ did not exhibit ATP dependence, displaying a dwell time of ~8 ms for all ATP concentrations. We concluded that ATP binds (presumably to the microtubule-bound leading head (*31*)) when kinesin is in its 1HB state and the unbound head is in-between previous and next microtubule binding sites.

## ATP hydrolyzes in two-head-bound state

Subsequently, we recorded traces of the kinesin construct T324C with 1 mM ATPγS, a slowly hydrolyzing ATP analogue. The results revealed that the use of ATPγS did not affect the unbound state duration, but drastically increased the time spent in the bound state (Fig. 4E). Therefore, we conjectured that ATP hydrolysis does not take place in the 1HB state, but after the unbound head moved to its next binding site. This hypothesis was further supported by comparing the run length (actual distance traveled) and run fraction (ratio of run length and distance from starting point to end of microtubule) determined from kymographs of Total Internal Reflection Fluorescence (TIRF) microscopy measurements for construct T324C under ATPγS and under ATP consumption. For 1 mM ATPγS, kinesin walked 36 times more slowly, but with an average run length nearly as long and an average run fraction nearly as large as for 1 mM ATP. (The observed slight reduction of run length is likely due to increased bleaching over the much longer run time.) Interestingly, while walking speeds were comparable for 1 mM ATPγS and 1 μM ATP, the run length was substantially shorter and the run fraction substantially smaller for 1 μM ATP (fig. S9). As the 1HB state is known to be the state that is most vulnerable to kinesin detachment from the microtubule (*35*), the combined results disqualify the 1HB state, i.e. the state in which a head is in-between the previous and next binding site on the microtubule, as the ATP-hydrolyzing state.

Further examination of the step successions in traces of construct T324C revealed a rare occurrence of an uneven number of 8-nm substeps between regular 16-nm steps (fig. S10). Since no concurrent side-stepping was observed, these substeps cannot be explained by the unbound labeled head docking to an adjacent microtubule protofilament at a binding site next to the leading head and kinesin subsequently switching protofilaments. We reasoned that the rare additional substeps probably arose from kinesin detaching into a weakly bound slip state which has so far only been reported for kinesin under load (*27, 36*) and not for freely walking kinesin. For cases in which the leading head was replaced by the trailing head on the same microtubule position, an uneven number of intermediate substeps is observed in the traces.

## Reconstruction of unbound head orientation

To explore the 3D-orientation of the unbound head, we repeated the 2D-MINFLUX tracking experiments with two additional kinesin constructs: E215C labeled at its solvent-exposed cysteine located at the C-terminal end of the β6-sheet (Fig. 5A top left) and K28C labeled at its solvent-exposed cysteine at aminoacid position 28 (Fig. 5A top right). In the bound state, the labeling positions E215C and K28C are located in the very front and back of the head, respectively, relative to the walking direction. Surprisingly, the fraction of detected pairs of substeps (B→U and U→B) of construct E215C did not differ substantially for 10 μM, 100 μM and 1 mM ATP. Unlike in previous experiments (Fig. 4C), these fractions always amounted to only ~10% of the entire population of substep pairs and regular steps (Fig. 5A bottom left). In fact, compared to construct T324C - especially for 10 μM ATP - this fraction was remarkably smaller and the unbound state was significantly shorter (fig. S11). In contrast, construct K28C (Fig. 5A right) did not notably differ in these aspects from construct T324C, indicating that the observed substeps of E215C do not represent the real 1HB state of the labeled head (for details, see supplementary text 2.4). Interestingly, the substep peak of the step size histogram of construct K28C was remarkably sharper than that of T324C, with a clear peak at 8 nm. Plotting the sequence of consecutive step sizes in 2D histograms revealed an underlying asymmetry of 6 nm B→U substeps and subsequent 10 nm U→B substeps of construct T324C, which is in stark contrast to the symmetric substepping behavior of the K28C construct (Fig. 5B).

**Fig. 5.**
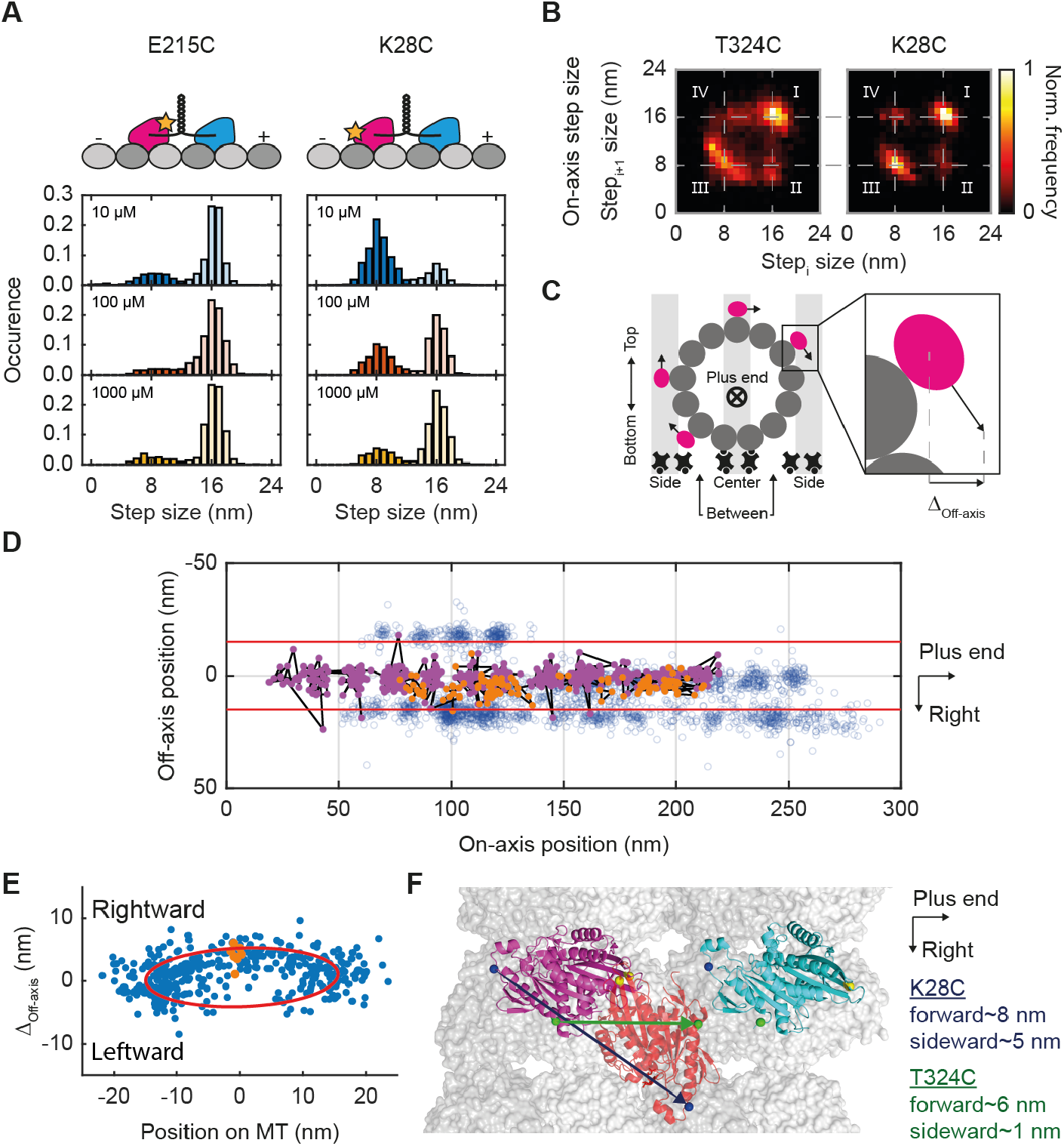
Rotation of the unbound head reconstructed through tracking various labeling sites on kinesin head. **(A)** Left: Construct E215C with fluorophore labeling site on the ‘front’ end of the kinesin head and histogram of detected step sizes of this construct for 10 μM, 100 μM and 1 mM ATP concentration featuring a dominant peak at 16 nm (N_10μM_=728, N_100μM_=380, N_1mM_=414) and a small fraction of 8 nm steps (N_10μM_=216, N_100μM_=65, N_1mM_=81). Right: Construct K28C with labeling position at the ‘back’ of the kinesin head and histogram of detected step sizes of this construct for 10 μM, 100 μM and 1 mM ATP concentration featuring an increasing peak at 16 nm (N_10μM_=186, N_100μM_=176, N_1mM_=601) and a decreasing fraction of 8 nm steps (N_10μM_=601, N_100μM_=130, N_1mM_=204) for increasing ATP concentrations. **(B)** 2D histograms of consecutive step sizes for constructs T324C and K28C both showing successive regular steps (I), regular steps followed by substeps (II), successive substeps (III) and substeps followed by regular steps (IV). For construct K28C, transitions involving the unbound state generate symmetric successive steps of around 8 nm. For construct T324C, these transitions exhibit an asymmetry of ~6 nm substeps that are followed by ~10 nm substeps. **(C)** Schematic of a surface-immobilized microtubule and the assignment of protofilament classes. The assumed true sideward displacement of the kinesin head (magenta) in these classes is indicated by black arrows. The zoom-in visualizes the projection of the three-dimensional displacement onto the imaging plane. **(D)** Scatter plot of four kinesin traces recorded on a single microtubule (blue circles) with one central trace highlighted for the detected bound (dark magenta dots) and unbound (orange dots) states. Red lines display the microtubule outline inferred from the traces. **(E)** Sideward displacement of all detected substeps and their respective position on the microtubule (blue dots). The orange dots correspond to the sideward displacement of the central trace shown in (E). The red line displays an ellipse fit to the data with major axis diameter of 30.6 nm and minor axis diameter of 9.2 nm. **(F)** Proposed 3D orientation of the unbound state of the trailing head (orange; PDB: 1MKJ) on the microtubule (alpha-tubulin in dark grey, beta-tubulin in light grey, PDB: 6DPU). Colored arrows depict the displacement of the different labeling positions from the microtubule-bound state of the trailing head (dark magenta; PDB: 3J8Y). The leading head is shown in the apo state (cyan; PDB: 4ATX).

We cannot fully exclude that the modification at amino acid position 215 hampered kinesin from entering a detectable 1HB state due to altered protein-protein interactions or steric hinderance. However, this scenario disagrees with earlier Förster-resonance-energy-transfer (FRET)-based observations of the 1HB state of a kinesin construct that was point-mutated and labeled at amino acid position 215 (*37*). Hence, we advance the following reasoning: As MINFLUX tracks the fluorophore position, our traces actually represent displacements of individual amino acids, more specifically, projections of the 3D amino acid trajectories onto the focal plane. Differences in substeps between individual kinesin constructs can be attributed to different trajectories of the amino acid positions 28 (aa28), 215 (aa215) and 324 (aa324) in space, pertaining to the ‘back’, ‘front’ and ‘mid’ part of the head, respectively.

Altogether, this allows us to reconstruct the 3D orientation of the labeled head in its unbound state. Under the assumption that each kinesin construct entered the 1HB state with its unbound head in-between previous and next microtubule binding sites, the non-apparent displacement of aa215 along the on-axis actually indicates a rotation of the unbound head around its front during a substep. Likewise, the symmetry of substep pairs of construct K28C implies a forward rotation of aa28 by about 8 nm, and the asymmetry of substep pairs of construct T324C suggests a displacement of aa324 by ~6 nm along the microtubule axis when entering the unbound state.

Next, we investigated the off-axis displacement of the unbound head which has been reported to be rightward (*16*) or non-existent (*15*) by different bead-tracking studies using an inherently artefact-prone label (*28*). We found off-axis displacements in the unbound state for constructs T324C and K28C (fig. S12). For construct T324C, they were small (<2 nm), whereas for K28C, off-axis displacements up to ~5 nm were measured. For the entire substep population of the latter, significant rightward, near-zero and leftward off-axis displacements appeared. Remarkably, the displacement was always consistent in magnitude and direction within a single trace as shown by Pearson correlation analysis (ρ=0.60±0.02).

To correlate the observed off-axis displacements with individual protofilament classes of a single microtubule (‘sides’, ‘center’, ‘between’) (Fig. 5C), we recorded trace sets of construct K28C on 19 different microtubules using actively stabilized samples (for details, see Materials and Methods section 1.1.3 and fig. S13). The outermost traces of the individual sets were spaced by ~30 nm, which agrees well with the microtubule outer diameter of ~25 nm (*38, 39*) plus twice the ~2.5 nm distance between the labeling position of construct K28C and the microtubule surface (inferred from PDB 3J8Y). Thus, these traces were assigned to kinesins walking along ‘side’ protofilaments and used as references for the remaining trace assignment.

Located between two ‘side’ traces, the ‘center’ traces nicely exhibit pronounced rightward displacements of the unbound states (Fig. 5D). After aligning all trace sets along their central on-axes, the substep off-axis displacements were plotted against the lateral offsets of the corresponding traces from the microtubule center axis (Fig. 5E). While imperfections in the alignment of all trace sets might have introduced minor errors in position, the traces of kinesins walking along ‘side’ protofilaments displayed a near-zero off-axis displacement in their substeps, ruling out a significant displacement of aa28 away from the microtubule surface. As protofilaments at the bottom of microtubules were mostly blocked by neutravidin and polymer (Fig. 5C), virtually all of the center traces can be attributed to kinesins walking on the microtubule top. Remarkably, these traces showed maximum and predominantly rightward substep off-axis displacements. This finding is corroborated by further detailed analysis of the ‘between’ traces, generated by kinesins able to walk both on ‘top’ and ‘bottom’ protofilaments, leading to both rightward (for ‘top’) and leftward (for ‘bottom’) off-axis displacements due to simple geometric reasons (see supplementary text 2.5). We conclude that upon entering the unbound intermediate state, aa28 of the trailing head is displaced up to ~5 nm rightwards with respect to the kinesin coordinate system.

The combination of on-axis substep sizes and associated off-axis displacements of the investigated amino acid positions allowed us to derive an approximate average 3D orientation of the unbound head during kinesin motion (Fig. 5F), improving the one derived from FRET studies of stationary kinesins(*37*). In conjunction with the coiled-coil rotation (Fig. 3), which is expected to resolve torsion and thus decrease asymmetry caused by a head’s full step, our data of systematic rightward displacement of the unbound head indicate that the hand-over-hand mechanism is symmetric.

## Mechanochemical cycle of kinesin-1

Combining all results, including reported observations, allowed us to refine the mechanochemical cycle of kinesin movement (Fig. 6). Starting with kinesin in its 2HB state, ATP in the trailing head hydrolyzes and phosphate (P_i_) exits the catalytic center (*40*). The trailing head detaches from the microtubule and, performing a substep, rotates into a rightward-displaced unbound state. Upon binding of ATP to the microtubule-bound leading head (*31*), the neck linker of the leading head partially docks (*41, 42*), moving the unbound head towards its next binding site and rotating the coiled-coil domain around its axis. The 16-nm step is completed after the (initially) trailing head releases adenosine-5’-diphosphate (ADP), binds to the microtubule and the (initially) leading head hydrolyzes ATP.

**Fig. 6.**
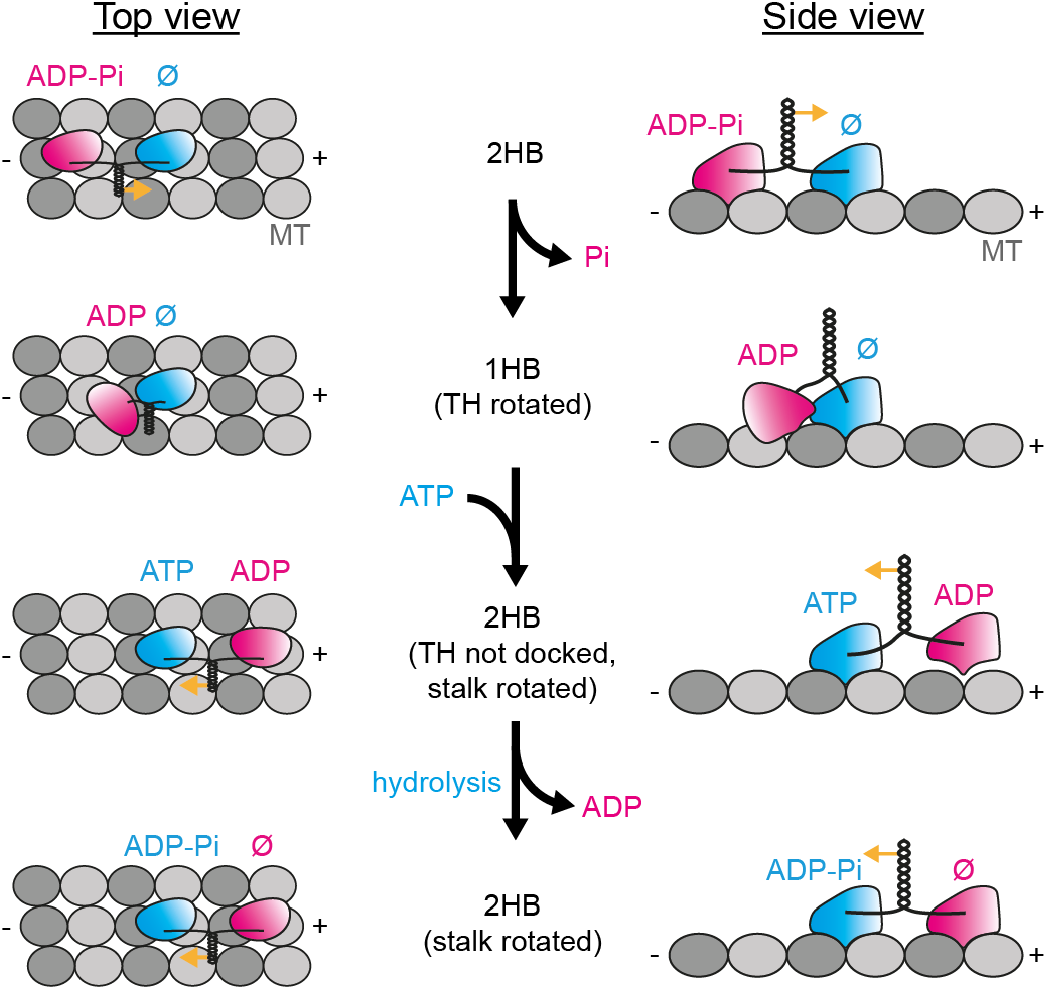
Head and coiled-coil rotation in the mechanochemical cycle of kinesin. Semi-cycle of kinesin walking along microtubules (MT) with 8 nm center-of-mass displacement in top and side view. Upon phosphate (P_i_) release*, the trailing head (TH, magenta) detaches from the MT and rotates into its rightward displaced unbound state. ATP binding to the leading head (LH, cyan) in the apo state (Ø) causes its neck linker to partially dock to the LH*, moving the TH towards its next binding site and significantly rotating the coiled-coil domain around its axis (as indicated by yellow arrows). The semi-cycle is completed by ATP hydrolyzing in the leading head* and the trailing head binding to the microtubule after ADP release*. (Aspects of the mechanochemical cycle that are inferred from literature are denoted by *.)

We note that our kinesin model is based on a fast MINFLUX tracking modality with 4 nm / 0.63 ms spatiotemporal resolution per individual localization using just 20 detected photons, allowing the identification of >12,000 kinesin steps with 1 nm precision (fig. S14). While this performance sets a new benchmark in the spatiotemporal tracking of proteins, further substantial improvements are possible in the future, for example by resorting to fluorophores with higher emission rates. Since it strongly suppresses background, the confocal arrangement of the MINFLUX system should also allow for detailed investigations of kinesin in living cells. Finally, our study clearly establishes MINFLUX as a next-generation tool for studying the dynamics of single proteins with minimal invasion. In fact, since nanometer MINFLUX tracking of single fluorophores requires less than a millisecond, one can readily envisage applying MINFLUX to any similarly fast nanometer-scale changes of fluorescently labeled biomolecules.

## Author contributions

J.O.W. and T.E. built the microscope, and together with J.E. wrote the software for controlling the setup and performing measurements. J.E. designed the interferometric system, with input from S.W.H., and provided technical supervision. J.O.W. built the active stabilization unit, wrote software for MINFLUX track evaluation and performed measurements, initially together with T.E.. L.S. scrutinized the kinesin tracking literature, identified, designed, prepared, and optimized the constructs for this study, advised by J.M. L.S. and J.O.W processed, evaluated and interpreted kinesin measurements, co-supervised by J.M., who was also responsible for the project administration. S.W.H. proposed and initiated the evaluation of MINFLUX spatio-temporal resolution for protein tracking and was responsible for overall supervision and steering. L.S., J.O.W., J.M. and S.W.H. wrote the manuscript with feedback from all authors. All authors contributed to the results through critical discussions throughout the course of the project.

## Acknowledgments

We thank Miroslaw Tarnawski and the Protein Core Facility at the MPI Heidelberg for performing the QuikChange site-directed mutagenesis and protein expression, our colleague Michael Remmel for access to the TIRF microscope, the MS Core Facility at the MPI Heidelberg for recording the mass spectroscopy data, Ron Vale (University of California, San Francisco and Howard Hughes Medical Institute) for providing plasmids K560CLM T324C and CLM RP HTR, Ahmet Yildiz (University of California, Berkeley) for providing plasmid K560CLM E215C as well as the protein preparation and purification protocol, and Matthias Kulp of the Optical Facility (MPI Göttingen) for imprinting aluminum gratings on coverslips.

