## Supplements for "MINFLUX dissects the unimpeded walking of kinesin-1"

#### Contents of Supplementary Materials

1 Materials and Methods ... 3

1.1 MINFLUX setup ... 3

1.1.1 Phase scanner ... 4

1.1.2 Phase scanner calibration ... 4

1.1.3 Active sample stabilization ... 5

1.2 TIRF microscope ... 6

1.3 Surface immobilized ATTO647N experiments ... 6

1.3.1 Flow chamber construction and surface coating ... 6

1.3.2 ATTO 647N surface immobilization and tracking buffer ... 6

1.3.3 Localization and tracking experiments ... 6

1.3.4 ATTO 647N data analysis ... 7

1.3.5 Step-finding algorithm ... 7

1.4 Kinesin experiments ... 8

1.4.1 Expression of kinesin 8
1.4.2 Labeling of kinesin 8
1.4.3 Preparation of microtubules 8
1.4.4 Sample preparation 9
1.4.5 Kinesin tracking experiments 9
1.4.6 Kinesin data analysis 9
1.4.7 Hidden Markov Model 10
1.4.8 Dwell time distribution model 10
1.4.9 TIRF imaging conditions and data analysis 11
1.5 Data representation 11
2 Supplementary Text 11
2.1 One-dimensional localization principle 11
2.2 Exponential gain in localization precision 12
2.3 Assignment of 16 nm steps 13
2.4 Artificial noise induced substeps 14
2.5 The unbound state's true displacement 14
3 Supplementary Figures 15
4 Supplementary Tables 29
Supplementary References 32

### 1 Materials and Methods

#### 1.1 MINFLUX setup

All position tracking experiments were performed on the custom-built MINFLUX confocal microscope schematically illustrated in Fig. S2. The light of two single frequency continuous-wave (cw) diode pumped excitation lasers with wavelengths of 642 nm (Bolero, Cobolt AB) and 488 nm (Calypso, Cobolt AB) are overlapped by dichroic mirrors and pass an acousto optical tunable filter (AOTF; 97-01776-0, EQPhotoncs) for power control before being coupled into a single mode fiber (P1-405BPM-FC-5, Thorlabs Inc.). A half-wave plate matches the light and fiber polarization angle. After the fiber, the polarization state is cleaned up again by a second half-wave plate and a Glan-Thompson-Polarizer to warrant a single polarization state and optimal power output. To optimize the depth of interference by varying the power balance between the subsequently generated beamlets, another half-wave plate is installed on a motorized rotation mount (PRM1, Thorlabs Inc.).

Next, the laser beam passes the interferometric phase scanner described in detail in supplementary section 1.1.1, which generates either two horizontally separated or two vertically separated beamlets. After the phase scanner, the polarizations of these four beamlets are adjusted to be orthogonal to the axis of displacement. As the opposing beamlets initially feature orthogonal polarization states, a custom-built segmented half-wave plate ensures alignment of these states.

The beamlets then pass a 90:10 (R:T) non-polarizing beam splitter cube (BS076, Thorlabs Inc.). The transmitted beamlets (10%) are split up twice by non-polarizing beam splitter cubes to be used for measuring the laser power and for calibrating the electro-optical phase modulator (EOPM) and the electro-optical amplitude modulator (EOAM) by monitoring the beamlets with CMOS cameras (Basler Ace acA1920-155µm, Basler AG) in a plane conjugated to the back focal plane (BFP) and in a plane conjugated to the focal plane (FP). Details of the phase scanner calibration are given in supplementary section 1.1.2.

The reflected beamlets (90%) are directed into the galvanometer scanner by a quad band dichroic mirror (ZET 405/488/561/640nm-TRFv2, Chroma Technology). The galvanometer scanner is a custom-designed implementation of a quad-scanner (*1*) consisting of four digitally encoded galvanometric scanning mirrors (GM-1015 with Ag mirrors, Canon Precision Optical Industry Co. Ltd.), which are steered by two controllers (GC-211, Canon). Each controller drives one x- and one y-mirror to ensure the best possible distribution of power load between both controllers. With a magnification of 100× with respect to the object space, the galvanometer scanner has a field-of-view (FOV) of 35×35 $\mu m^2$ with a resolution given by the digital encoders of less than 1 nm. The positional jitter produced by the feedback loop is below 1 nm.

After the galvanometer scanner, the beamlets are coupled into the rear infinity port of a Leica DMi8 microscope body and guided by a mirror in the filter wheel towards the objective lens (Leica HC PL Apo 1.4 100x OIL STED white, Leica Microsystems). The microscope body further offers options for fluorescence widefield illumination, observation via eyepiece or camera detection. The sample is mounted on a custom-built glass stage that can be moved in three dimensions (3D) over the range of several millimeters by piezo actuators with 1 nm precision encoders (SLC1730, Smaract GmbH).

Fluorescence light emitted from the sample travels back through the microscope body, is descanned by the galvanometer scanner and is decoupled from the excitation path at the quad band filter. Afterwards, it is focused onto a motorized pinhole (MPH16-A. Thorlabs Inc.) by a 400 mm achromatic lens leaving a total magnification of 200x. The pinhole can be adjusted from 25 μm to 2 mm, which translates into 125 nm to 10 μm in the object space. Typically, the pinhole size was set to 350 nm in object space, which is slightly larger than the periodicity of the excitation interference pattern in the FP. Fluorescence light passing the pinhole is collimated and guided towards one of two spectrally separated avalanche photo diode (APD) detectors (SPCM-AQRH-13-ND, Excelitas Technologies Corp.) by a cascade of dichroic filters and detected between 500 nm and 550 nm ('green'; 525/50 nm BrightLine single-bandpass filter, Semrock) and between 662 nm and 800 nm ('red'; 732/137 nm BrightLine single-bandpass filter, Semrock). To ensure drift-free, long-term measurements, the MINFLUX setup actively stabilizes the sample position by imaging an aluminum grating imprinted onto the coverslip with an infra-red laser coupled into the microscope body. A detailed description of the active stabilization unit is given in supplementary section 1.1.3.

The excitation laser light that is reflected from the sample also travels back through the microscope body, is reflected at the quad band filter, passes the 90:10 (R:T) non-polarizing beam splitter cube and is then focused on a separate 25 μm pinhole before a photon multiplier tube (PMT; H14119-40, Hamamatsu Photonics). This signal was mainly used to find and adjust the axial position of the sample to the focal point of the objective lens, but can also be used to gain information about local scattering or reflection processes.

To ensure an exact timing of the fast measurement process, the data acquisition, relevant processing and hardware control is executed by a field programmable gate array board (FPGA; PCIe-R7852r, National Instruments). The FPGA board employs a 100 MHz clock, several analog outputs (AO) with 16 bit precision and a 1 MHz update rate, several analog inputs as well as digital inputs and outputs (DIO). Programming of the FPGA was performed in LabView 2019 (National Instruments). During measurements, the FPGA controls the galvanometer scanner via DIO, the phase scanner via AO, records detected photons via DIO and AI, computes the estimated molecule position and communicates with the host computer via first-in-first-out (FIFO) memories.

##### 1.1.1 Phase scanner

The interferometric phase scanner (Fig. S3) is implemented to generate excitation intensity distributions with line-shaped minima ($LS_x$ and $LS_y$) and move their minima along the x- and y-direction, respectively. To this end, a single, horizontally polarized laser beam passes the EOPM consisting of two orthogonal electro-optical crystals (RTP, custom manufactured by Cristal Laser, P.A. du Breuil, France) aligned at a 45° angle with respect to the optical table. These crystals impose a phase difference $\Delta\phi$ between the two polarization components along the crystal axes depending on the applied voltage. The two components are then separated by a polarizing beam displacer (BD) aligned at a 45° angle. The resulting two beamlets pass the EOAM consisting of a pair of electro-optical crystals (RTP, Cristal Laser) aligned at a 0° angle. The crystals are used to either keep the incoming polarization state or rotate it by 90°. Afterwards, the beamlets pass another polarizing BD oriented 90° to the first one, thereby generating either two horizontally separated or two vertically separated beams. Since opposite beams are orthogonally polarized, a segmented $\lambda/2$-plate is used to align these polarizations.

##### 1.1.2 Phase scanner calibration

The phase scanner requires information on the volt-to-nanometer scaling of the EOPM as well as on the voltages generating the $LS_x$ and $LS_y$ excitation intensity distribution. To calibrate these values, in the first step, the voltage applied to the EOAM was varied while simultaneously recording images with the BFP-camera. The resulting image stack was analyzed by identifying the four spots and tracking their intensity with respect to the applied voltage. The voltages that simply rotate the input polarization without adding ellipticity generated dips in the signal, which in turn were fitted with a parabola (Fig. S3). These dips identified the voltages that generated a single pair of beamlets and created the $LS_x$ and $LS_y$ excitation intensity distribution. To increase contrast, the laser power was increased such that the beamlets typically saturated the camera.

In the second step, the EOAM was set to create the $LS_x$ excitation intensity distribution on the FP-camera and the voltage applied to the EOPM was modulated, while recording images with the FP-camera. The resulting image stack was analyzed by identifying the focal distribution and tracking its modulation at the central pixel. The modulation was fit with a sine function to extract the voltage-to-phase scaling and the phase shift (Fig. S3). Repeating the same procedure for the $LS_y$ excitation intensity distribution yielded the corresponding parameters in y-direction.

Finally, single surface-immobilized fluorescent nanospheres (F8782, Thermo Fisher) were tracked while moving them with the piezo-electric stage by a defined distance. As the sample position scales linearly with the phase, the data were used to calibrate the phase-to-nanometer scaling.

##### 1.1.3 Active sample stabilization

To actively stabilize the sample with high precision and without adding fiducial makers to it, we used coverslips coated with a thin (~3 nm) aluminum grating imprinted onto the coverslip side facing the objective lens. This grid was imaged by two oppositely tilted infra-red laser beams in widefield configuration (Fig. S13), which are generated by deflecting two segments of a 830 nm collimated laser diode (CPS830S, Thorlabs Inc.) with a rooftop shaped glass plate and coupling them into the microscope body below the objective lens with a short-pass filter (T790spxrxt-UF3, AHF Analysetechnik). Light reflected from the grid was imaged onto a CMOS camera (Basler Ace acA4112-20µm, Basler AG) generating two spatially separated image segments corresponding to the two illuminating beams. The mean lateral drift of the two image segments accounted for lateral drift of the grid pattern on the camera and consequently for lateral sample drift. Due to the opposite tilt of the beams, axial sample drift could also be directly measured as the difference in drift between the image segments.

The camera recorded frames at 20 fps, which were accumulated for 500 ms before further processing. First, the accumulated image was separated into two segments corresponding to the two illuminating beams and simultaneously rotated such that the grid was aligned with the camera axes. Next, the mean x- and y-profiles of each segment were calculated, Fourier-transformed, and the main frequency peak representing the grid periodicity was identified. The phase of this peak was used to calculate the 'position' of the grid for each image segment individually. This position was rotated back into the camera coordinate system and used to calculate the corresponding 3D- drift on the camera

$$\begin{pmatrix} x_{cam} \\ y_{cam} \\ z_{cam} \end{pmatrix} = \begin{pmatrix} x_1 + x_2 \\ y_1 + y_2 \\ y_1 - y_2 \end{pmatrix} /2 \,. \qquad (1)$$

To translate this into the actual drift of the sample, we introduced a calibration matrix $M_{3x3}$ that accounted for the magnification between sample and camera plane as well as for imperfections of the camera-to-stage alignment

$$\begin{pmatrix} x_{sample} \\ y_{sample} \\ z_{sample} \end{pmatrix} = M_{3x3} \begin{pmatrix} x_{cam} \\ y_{cam} \\ z_{cam} \end{pmatrix}. \tag{2}$$

#### 1.2 TIRF microscope

Total internal reflection fluorescence (TIRF) measurements were performed on a custom-built widefield setup equipped with a 473 nm ('green') excitation solid state cw laser (gem 473, Novanta) and 642 nm ('red') excitation cw fiber laser (MPB Communications Inc.), a back-illuminated EMCCD camera (iXon 897 / 512×512 sensor, Andor) and a 100x/1.46 NA oil immersion lens (HCX PL APO CS, Leica). Fluorescence emission was separated from the excitation light using dichroic mirrors ('green': T505lpxr, Chroma; 'red': HC 660, Semrock) and detected between 560 nm and 594 nm ('green'; 550/88 nm BrightLine single-bandpass filter, Semrock) and between 664 nm and 736 nm ('red'; ET700/75 nm, Chroma). TIRF illumination mode was achieved with a movable mirror.

#### 1.3 Surface immobilized fluorophore (ATTO647N) experiments

##### 1.3.1 Flow chamber construction and surface coating

Flow chambers were constructed by attaching oxygen-plasma-cleaned coverslips to objective slides with double-sided adhesive tape. The chambers were incubated with 0.2 mg/ml biotinylated poly-L-lysine-polyethylene-glycol (PLL-PEG-bt) solution (PLL(20)-g[3.5]-PEG(2)/PEG(3.4)-biotin, Susos AG Inc.) supplemented with 1 % (v/v) Tween 20 (P9416, Sigma-Aldrich) in dd$H_2O$ for 15 min, rinsed with PEM80, incubated with 10 µg/ml neutravidin (NVD; 31000, Thermo Fisher) in PEM80 for 5 min, and rinsed with PEM80.

##### 1.3.2 ATTO 647N surface immobilization and tracking buffer

Biotinylated ATTO 647N (stock solution 1 mM in dimethyl sulfoxide (DMSO); *AD 647N-71, ATTO-TEC*) was diluted in PBS to single molecule conditions of 30 fM, flushed into the flow chambers and incubated for 5 min. After rinsing with PBS, imaging buffer (1 mM methyl viologen (75365-73-0, Sigma Aldrich), 1 mM ascorbic acid (50-81-7, BioVision Inc.) in PBS) was flushed into the flow chambers and the chambers were sealed with picodent twinsil speed 22 (picodent Dental-Produktions- und Vertriebs GmbH).

##### 1.3.3 Localization and tracking experiments

All position tracking experiments were performed on the custom-built MINFLUX confocal microscope described in supplementary section 1.1. For tracking surface-immobilized ATTO 647N molecules, confocal xy-scans were recorded to identify the diffraction-limited positions of molecules within a 10×10 µm$^2$ FOV. These positions were then individually addressed by the galvanometer scanner to run MINFLUX tracking measurements. During the initial zoom-in process, up to five MINFLUX iterations with decreasing $L$ were performed to bring the excitation intensity minimum close to the actual molecule position. Until photobleaching, all further localizations were performed with the smallest $L$ and the excitation intensity minimum was in each dimension repeatedly addressed to positions at [-$L$/2, 0, $L$/2] centered on the latest molecule position estimate. After each cycle, the molecule position estimate was updated based on the detected photon counts. The exposure time for each of the three exposures per dimension was set to 100 µs. Together with 5 µs dead time to switch between exposures and 500 ps for calculating the updated molecule position, a single two-dimensional (2D) localization took 631 μs, which corresponds to a sampling rate of around 1.5 kHz. Detailed information on all MINFLUX routines is given in Table S3.

##### 1.3.4 Fluorophore (ATTO 647N) data analysis

MINFLUX position traces of surface-immobilized ATTO 647N molecules were individually screened and filtered in LabView 2019 by a minimum of five photons per localization in each dimension, a maximum of 60 photons per localization in each dimension, and a minimum number of ten localizations. The molecule position estimate was then recalculated using a fixed-curvature estimator, which is more efficient for low photon numbers. As this estimator requires a robust estimate of the curvature of the parabola approximating the excitation intensity minimum, the curvature $c$ was estimated from all localizations of a single trace using

$$c = \frac{2}{L^2}(n_+ + n_- - 2n_0)\,, \tag{3}$$

with $n_-$, $n_0$ and $n_+$ being the photon counts detected during exposures at positions -$L$/2, 0 and $L$/2. For $c$<0, positions were discarded. Position traces with a good average signal-to-background ratio (SBR) were manually selected and exported to be further analyzed with dedicated scripts in MATLAB. The average localization precision $\sigma$ a trace with length $K + 1$ was estimated by collectively fitting a Gaussian function to the distribution of pairwise distances $x_{i+1} - x_i$ in both dimensions and by dividing the obtained standard deviation by $\sqrt{2}$

$$\sigma = \sqrt{\frac{1}{2K}\sum_{i=1}^{K}(x_{i+1} - x_i)^2}\,. \tag{4}$$

The $SBR$ was estimated from the mean photon counts of the three exposures using

$$SBR = \frac{n_+ + n_-}{2n_0} - 1\,. \tag{5}$$

##### 1.3.5 Step-finding algorithm

To detect steps in individual MINFLUX position traces featuring a localization precision $\sigma$, the MATLAB function *ischange* was applied that uses an iterative change point search minimizing

$$\sum_{i=1}^{m+1}\left[C\left(y_{\tau_{i-1}+1:\tau_i}\right)\right] + \beta m\,, \tag{6}$$

with $C$ being a cost function for individual segments $i$, and $\beta$ being a penalty for adding single steps to prevent overfitting. Typically, the cost function is based on the maximum likelihood estimate for finding individual change points, which is equivalent to the residual sum of squares (RSS). As adding steps to traces with a constant mean results in a RSS reduction close to $\sigma^2$, the overfitting rate of simulated data for different noise levels was numerically evaluated by varying the penalty factor between $\sigma^2$ and $20\sigma^2$. Since the simulations did not show any dependence on the localization precision level, requiring <0.01% overfitting (1 artificial step per 10,000 localizations) yields an optimal penalty factor of $12\sigma^2$ that can be applied independently of the localization precision. The localization precision of individual traces was estimated as described for the surface-immobilized ATTO 647N data (see supplementary section 1.3.4, eq. (4)) and thus can be evaluated without any prior knowledge about steps.

After running the step-finding algorithm, two filters were applied to the detected step function. The first filter removed steps originating from spikes in the data by using a moving median filter of width nine, thus eliminating spikes of width four. Note that this filter does not affect short steps which occur unidirectionally in succession. The second filter removed steps of step sizes smaller than a certain threshold. This threshold was set to 5 nm for tracking data of the kinesin-1 head and to 2 nm for tracking data of the coiled-coil domain.

The step detection probability was numerically simulated for different step-size-to-noise ratios and varying plateau durations (Fig. S6) by generating traces with localization precision $\sigma$ and adding two steps of step size 8 nm spaced by $1{:}n$-data points. For a typical localization precision of 4 nm, the step-finding algorithm showed a reduced step detection efficiency for the simulated unbound state shorter than 8 ms.

#### 1.4 Kinesin experiments

##### 1.4.1 Expression of kinesin

Cysteine-light truncated (at amino acid position 560) human kinesin-1 constructs (hereafter referred to as kinesin) were expressed in E. coli using the plasmids K560CLM T324C (#24460, Addgene) (*2*), K560CLM E215C (kindly provided by the Yildiz Lab, University of California, Berkeley), and K560CLM K28C and K560CLM N356C both produced from CLM RP HTR (#24430, Addgene) (*3*) by QuikChange site-directed mutagenesis (using PfuUltra HF polymerase (600380-51), Agilent) as described by Tomishige et al. (*3*). All plasmids were verified by DNA sequencing. The constructs each contained a single solvent-exposed cysteine at amino acid position 324, 215, 28 or 356 for labeling, and a C-terminal $His_6$-Tag for purification (via 5 ml HisTrap FF (GE17-5255-01, Cytiva)). The purified protein (in 25 mM piperazine-N,N′-bis(2-ethanesulfonic acid) (PIPES, P-1851, Sigma-Aldrich) pH 6.8, 2 mM $MgCl_2$ (1.05833.0250, Merck), 1 mM ethylene glycol-bis(2-aminoethylether)-N,N,N′,N′-tetraacetic acid (EGTA, E3889, Sigma-Aldrich), 0.1 mM ATP (BP413-25, Fisher Scientific), 0.2 mM TCEP (J60316, Alfa Aesar), around 300 mM NaCl (1.06404.1000, Sigma-Aldrich), 10% (m/v) sucrose (S1888, Sigma-Aldrich)) was aliquoted, flash frozen in liquid nitrogen and subsequently stored at -80°C.

##### 1.4.2 Labeling of kinesin

Kinesin was labeled with ATTO 647N maleimide (AD 647N-41, ATTO-TEC) over night at 4°C. Excess dye was removed from the reaction mixture by size-exclusion chromatography (PD MiniTrap G-25, 28-9180-07, Cytiva) according to the manufacturer's protocol. The degree of labeling (DOL) was determined by UV-Vis spectroscopy (DS-11+ Spectrophotometer, DeNovix) and mass spectrometry (ESI, maXis II ETD, Bruker) (Fig. S8). Sucrose was added to the labeled protein in a concentration of 10 % (w/v) and aliquots were flash-frozen in liquid nitrogen and stored at -80°C.

##### 1.4.3 Preparation of microtubules

Biotinylated and fluorescently labeled microtubules were polymerized from 88% Cycled Tubulin (032005, PurSolutions, LLC), 10% Labeled Tubulin-Biotin-XX (033305, PurSolutions, LLC) and 2% Labeled Tubulin-Alexa Fluor 488 (048805, PurSolutions, LLC). The lyophilized tubulin variants were suspended in PEM80 buffer (80 mM PIPES, 0.5 mM EGTA, 2 mM $MgCl_2$, pH 7.4) with 1 mM guanosine-5'-[(α,β)-methyleno]triphosphate (GMPCPP; NU-405S, Jena Bioscience) and the solution was incubated for 30 min at 37°C. Afterwards, the polymerized microtubules were centrifuged at 21,000x g in a bench-top microcentrifuge (Fresco 21, Thermo Scientific) for 15 min, washed with PEM80 and centrifuged at 21,000 × g for 15 min. The microtubule pellet was resuspended in PEM80, aliquoted, flash-frozen in liquid nitrogen and stored at -80°C.

##### 1.4.4 Sample preparation

The flow chamber described in supplementary section 1.3.1 incubated with microtubules diluted in 10 ug/ml cabazitaxel (FC19621, Biosynth Carbosynth) in PEM80 for 5 min, rinsed with PEM80, blocked with 100 µg/ml biotinylated bovine serum albumin (BSA-bt; A8549-10MG, Sigma-Aldrich) in PEM80 for 5 min, and rinsed with PM15 buffer (15 mM PIPES, 2 mM $MgCl_2$, pH 7.4). Labeled kinesin in measuring buffer (1 mM 1,4-dithiothreitol (DTT, 6908.1, Carl Roth), 1 mM paclitaxel (10-2095, Focus Biomolecules), 10 µg/ml BSA-bt, 1 mM methyl viologen, 1 mM ascorbic acid, 10 µM/100 µM/1 mM adenosine 5′-triphosphate (ATP; A3377-1G, Sigma-Aldrich) or 1 mM ATPγS (NU-406-5, Jena Bioscience) in PM15 buffer) was added in an Invivo2 Plus Hypoxia Workstation (0.3% O2, 0.1% CO2, 99.6% N2; Baker Ruskinn) and sealed with picodent twinsil speed 22 or nail polish.

##### 1.4.5 Kinesin tracking experiments

To record MINFLUX position traces of kinesin, individual microtubules were manually identified from 5×5 $\mu m^2$ confocal xy-scans with a pixel size of 50 nm using the 488 nm laser. Up to three pixels on a microtubule were selected and sequentially multiplexed with 10 ms exposures of the 640 nm laser. If the detected photon rate surpassed a threshold of 5 kHz, the MINFLUX tracking routine (as explained in supplementary section 1.3.3) was triggered. Detailed information on all MINFLUX routines is given in Table S3. For kinesin tracking experiments with ATP$\gamma$S, the triggering routine was adapted to compensate for the slow kinesin movement. A 10×10 $\mu m^2$ dual-color confocal xy-scan was recorded and emission spots in the kinesin channel that colocalized with microtubules were individually addressed for MINFLUX tracking measurements. Kinesin traces were recorded for about 1 s at high ATP concentrations, and up to 3 s for lower ATP concentrations and for ATP$\gamma$S.

##### 1.4.6 Kinesin data analysis

MINFLUX position traces of kinesin were individually screened and filtered adjusting the minimal number of photons required for a successful localization to exceed the number of background photons under the given measurement conditions. Traces displaying a clear stepping behavior and a good average SBR were manually selected, exported and subjected to a detailed analysis using dedicated MATLAB scripts. For each kinesin trace, the start and end point of analysis as well as the approximate position of the first kinesin step within this region were manually selected. Next, the average plateau position before the first step was subtracted from the entire trace in both dimensions and the corrected positions were used to align the linear movement of kinesin with the x-direction by a simple rotation. Step detection was performed along this direction with the step-finding algorithm described in supplementary section 1.3.5 to identify change points, i.e. kinesin steps. The same change points were applied in y-direction to extract the off-axis displacement. To determine for each plateau whether the labeled kinesin head was in its bound ($B$) or unbound ($U$) state, the sequence of detected step sizes was analyzed by a Hidden Markov Model (HMM) explained in supplementary section 1.4.7.

For evaluation of the coiled-coil rotation and the unbound head rotation, the data was pooled for individual ATP concentrations per kinesin construct and grouped by either the state or the state transition as derived from the HMM.

##### 1.4.7 Hidden Markov Model

To extract the most likely sequence of bound ($B$) and unbound ($U$) states of the labeled kinesin head, a five state HMM was constructed (Table S1) based on the sequence of detected step sizes as identified by the step-finding algorithm (see supplementary section 1.3.5). Main transitions are from $B$ to $B$, from $B$ to $U$, and from $U$ back to $B$ as these are most likely to occur. Emission values for these transitions are normally distributed:

$$E = \begin{cases} n * 16\,nm & B \to B \\ (2n+1) * 8\,nm & B \to U \\ (2n+1) * 8\,nm & U \to B \end{cases} \tag{7}$$

The transition matrix was designed such that the head cannot switch states between steps. To account for presumably rarely occurring $U$ to $U$ transitions as well as potential slip states, two additional transitions were included in the model:

$$E = \begin{cases} (2n+1) * 8nm & B \to B \\ n * 16nm & U \to U \end{cases} \tag{8}$$

The probability for these transitions was set to 1%, as bound states are significantly less likely to be missed than unbound states (see supplementary text 2.3) and assuming slip states to also only occur rarely. An overview of the assignment for the kinesin data is given in Table S2.

##### 1.4.8 Dwell time distribution model

To extract the average residence time kinesin spends in its one-head-bound (1HB) and in its two-head-bound (2HB) state, the dwell times of the individual $U$ and $B$ plateaus were evaluated and plotted in histograms. Any steps associated with back-steps were excluded from the temporal analysis as we focused on the mechanochemics of the processive forward motion. To account for the reduced step detection probability for steps of step sizes around 8 nm into plateaus shorter than 8 ms, only plateaus longer than 8 ms were included in the temporal analysis. Both histograms were fitted simultaneously using a single exponential decay $p_U$ with the time constant $\tau_{1HB}$ for the duration of the $U$ state

$$p_U(t) = \frac{e^{-\frac{t}{\tau_{1HB}}}}{\tau_{1HB}} \tag{9}$$

and a convolution of three exponential decays $p_B$ with the time constants $\tau_{1HB}$ and twice $\tau_{2HB}$ for the duration of the $B$ state

$$\begin{aligned} p_B(t) &= \int_0^{\tau} p_{2HB}(\tau - t) * \int_0^{t} p_{1HB}(t - t') * p_{2HB}(t')dt'\,dt \\ &= \frac{1}{\tau_{2HB}(\tau_{2HB} - \tau_{1HB})}\left(\tau e^{-\frac{\tau}{\tau_{2HB}}} - \frac{\tau_{1HB}\tau_{2HB}}{\tau_{2HB} - \tau_{1HB}}\left(e^{-\frac{\tau}{\tau_{2HB}}} - e^{-\frac{\tau}{\tau_{1HB}}}\right)\right). \end{aligned} \tag{10}$$

##### 1.4.9 TIRF imaging conditions and data analysis

For kinesin tracking at 1 µM ATP and at 1 mM ATPγS, the exposure time per frame and the 642 nm laser power were set to 10 s and 10 µW, and for measurements at 1 mM ATP to 100 ms and 300 µW. Imaged with these parameters, single kinesins (construct T324C) labelled with ATTO 647N immobilized on microtubules using the non-hydrolyzing ATP analogue adenylyl-imidodiphosphate (AMP-PNP, A2647-5MG, Sigma Aldrich) did not significantly bleach over average run times. Microtubules were recorded at 2 µW 473 nm laser power. All laser powers were measured in the back focal plane of the objective. Frame series were analyzed with the KymographBuilder in Fiji to determine the run length along microtubules (*4, 5*).

#### 1.5 Data representation

**Traces** illustrate the MINFLUX tracked fluorophore positions either over time or as xy-scatter plots together with their corresponding step function.

**Histograms** show the counts per individual bin normalized to the size of the population. Single datasets are displayed as bars and multiple datasets in a single plot as stair functions or lines.

**2D histograms** present the normalized occurrence of successive step size pairs for either on-axis displacement or for the off-axis displacement of consecutive $B \rightarrow U$ and $U \rightarrow B$ transitions.

**Cumulative density plots** are generated by counting the fraction of elements below the corresponding x-axis values.

### 2 Supplementary Text

#### 2.1 One-dimensional localization principle

Localizing along a single axis with a one-dimensional (1D) intensity minimum simplifies the mathematical analysis needed for on-the-fly position tracking. The molecule position is probed with an approximately parabolic excitation profile $y = b + c(x - x_M)^2$ at positions $-L/2$, 0 and $L/2$ relative to the previous position estimate $x_M$. The detected photon counts $n_-$, $n_0$ and $n_+$ can thus be described by the following set of equations:

$$n_- = b + c\left(-\frac{L}{2} - x_M\right)^2 + \epsilon_- \,, \tag{11}$$

$$n_0 = b + cx_M^2 + \epsilon_0 \,, \tag{12}$$

$$n_+ = b + c\left(\frac{L}{2} - x_M\right)^2 + \epsilon_+ \,, \tag{13}$$

with $\epsilon$ being the Poisson noise term. As this set is lacking information for being completely solved in a simple approximation, the noise terms are dropped and it is assumed that the measured photon counts match the average photons at the respective excitation intensities. The resulting equation for the molecule position estimate is then given by solving this set of equations analytically

$$x_M = \frac{L}{4}\frac{n_- - n_+}{n_- + n_+ - 2n_0}. \tag{14}$$

By only requiring knowledge about the probing distance $L$ without using further information on the exact excitation profile, eq. (14) thus does not need to be adapted to varying background contributions or excitation wavelengths. Moreover, this calculation can be directly implemented into a FPGA board for precise molecule position estimates.

In the following, the influence of the previously neglected noise terms on the molecule position estimate is considered. As the noise is assumed to be Poissonian, the variance of the individual photon counts is equal to their mean. By performing a simple Gaussian error propagation, the standard deviation $\sigma_x$ of the position estimate, i.e. the localization precision, is given by

$$\sigma_x = \frac{L}{2}\frac{\sqrt{(n_+ - n_-)^2 n_0 + (n_0 - n_-)^2 n_+ + (n_+ - n_0)^2 n_-}}{(n_+ + n_- - 2n_0)^2}. \tag{15}$$

As $n_-$, $n_0$ and $n_+$ represent the average detected photon counts, they can be substituted by the parabolic excitation profile. This shall only be done here for the case $x_M = 0$, as during tracking measurements the excitation intensity minimum is adjusted after each recorded localization, which always keeps the minimum close to the molecule. By further demanding for a constant number of photons, the localization precision is given by

$$\sigma_x = \frac{L}{4\sqrt{N}}\sqrt{1 + \frac{5}{2}\frac{4b}{cL^2} + \frac{3}{2}\left(\frac{4b}{cL^2}\right)^2} = \frac{L}{4\sqrt{N}}\sqrt{1 + \frac{5}{2SBR(L)} + \frac{3}{2SBR(L)^2}}. \tag{16}$$

As the term $cL^2/4b$ represents the ratio of the background corrected intensity at positon $\pm L/2$ relative to the background level $b$ in the excitation intensity minimum, we define this value as the signal-to-background ratio $SBR$. At large $L$, the influence of the $SBR$ on the localization precision can be neglected entailing a linear scaling with $L$. However, with decreasing $L$, the $SBR$ contribution becomes dominant, setting a lower limit to the achievable localization precision, which roughly scales with the inverse square root of the curvature-to-offset ratio of the parabola

$$\sigma_{x.min} \approx \sqrt{\frac{b}{c}\frac{1}{\sqrt{N}}}. \tag{17}$$

Setting aside increasing the molecule's emission rate, optimizing the STR of MINFLUX directly translates into minimizing the background $b$ and maximizing the curvature $c$ of the excitation profile.

#### 2.2 Exponential gain in localization precision

The iterative MINFLUX zoom-in process can be implemented in a self-contained manner by choosing the $L$ of the next zoom-in step $j$ based on the localization precision $\sigma$ of the previous step

$$L_j = \alpha\sigma_{j-1}. \tag{18}$$

Here, we denote $\alpha$ as the regulation parameter. It defines how strongly the zoom-in process is pushed and thereby ultimately defines the success rate of the algorithm. The smaller $\alpha$ is chosen, the faster the process culminates and the fewer photons are consumed, but the higher the probability is for a molecule to be outside of the probed region in the next zoom-in step. Now assuming a background free localization, the localization precision is obtained from eq. 16

$$\sigma_j = \frac{L_j}{4\sqrt{N_j}}. \tag{19}$$

Inserting eq. 18 into eq. 19 and zooming-in with $k$ steps yields a final localization precision of

$$\sigma_k = \frac{\left(\frac{\alpha}{4}\right)^k \sigma_0}{\sqrt{\prod_{i=1}^{k} N_i}}. \tag{20}$$

Here, the first probing distance $L_1 = \alpha\sigma_0$ is set relative to the initial localization precision $\sigma_0$, which does not originate from the MINFLUX measurement, but from the diffraction-limited pre-localization.

To achieve the best possible localization precision, the detected photon number of all individual zooming-in steps $i$ should be equal $N_i = N/k$, as this provides the highest product $\prod_{i=1}^{k} N_i$ with a fixed number of total detected photons *N*. By inserting this constraint into eq. 20 and performing the derivative for the number of zoom-in steps, the optimal number of steps $k_{opt}$, into which the zoom-in process should be split, can be calculated to be

$$k_{opt} = \frac{16N}{e\alpha^2}, \tag{21}$$

and combined with eq. 20 to yield the final localization precision

$$\sigma_k = \sigma_0 e^{-\frac{8N}{e\alpha^2}}. \tag{22}$$

We highlight that by bringing the excitation intensity minimum iteratively closer to the molecule position, the localization precision exponentially scales with the number of detected photons, making MINFLUX fundamentally more photon efficient than non-iterative processes. The only parameters that have to be independently optimized are the initial localization precision $\sigma_0$, providing the first rough molecule position estimate, and the regulation parameter $\alpha$, determining the push of the iterative MINFLUX zoom-in process.

#### 2.3 Assignment of 16 nm steps

As the HMM includes 16 nm steps for both $B$ to $B$ and $U$ to $U$ transitions, the ratio between these two transitions was estimated to allow for a distinctive assignment of states. To simplify the calculation, we assumed equal dwell times for the 1HB and 2HB state ($\tau_{1HB} = \tau_{2HB} = \tau$ = 8 ms). Under this constrain, the probability distribution for the dwell time $t$ of the bound state $B$ is given by

$$p_B(t) = \frac{t^2 e^{-\frac{t}{\tau}}}{2\tau^3}, \tag{23}$$

whereas the probability distribution for the unbound state is given by eq. (9). Combining the assumptions $\sigma$ = 4 nm and step size=8 nm with the numerically derived step detection probability $p_D$ (Fig. S6), we can calculate the ratio of missed substeps as follows:

$B \rightarrow B$: $$1 - \int p_U(t) * p_D(t)dt = p_{B\rightarrow B} \tag{24}$$

$$U \rightarrow U: \qquad 1 - \int p_B(t) * p_D(t) dt = p_{U\rightarrow U} \tag{25}$$

Additionally including information on not missing the states before and after the 16 nm steps results in an even stronger reduction of the probability of observing $U$ to $U$ transitions:

$$P_{BB} = (1 - p_{U\rightarrow U}) * p_{B\rightarrow B} * (1 - p_{U\rightarrow U}) \tag{26}$$

$$P_{UU} = (1 - p_{B\rightarrow B}) * p_{U\rightarrow U} * (1 - p_{B\rightarrow B}). \tag{27}$$

The resulting ratio of $B$ to $B$ and $U$ to $U$ transitions is then given by

$$\frac{P_{B\rightarrow B}}{P_{U\rightarrow U}} \approx 51\,. \tag{28}$$

#### 2.4 Artificial noise-induced substeps

The step-finding algorithm, which was used to analyze the stepping motion of kinesin was optimized such that overfitting the MINFLUX tracking data was highly unlikely for a purely Gaussian position noise. However, when deviating from Gaussian noise, the addition of artificial steps to the detected step function could not be entirely excluded. This is expected to mainly play a role for data-points close to regular kinesin steps. Otherwise, either single steps with very small step sizes or paired steps, leaving and returning to the previous plateau, would appear. The former ones were removed during trace filtering (see section 1.3.5), while the latter ones were excluded as backsteps from further analysis. Artificial steps near regular steps would have caused these to split into two pseudo substeps and would have generated an artificial unbound state. Yet we expect the dwell time of this artificial unbound state to be short, since we consider longer deviations from Gaussian noise to be very unlikely.

#### 2.5 The unbound state's true displacement

As we do not assume any preferred protofilaments for kinesin to walk on, the only restriction is given by the 'bottom' filaments being blocked by neutravidin and polymer. Therefore traces in 'center' protofilaments solely represent kinesin walking on 'top' of the microtubule. 'Side' and 'between' traces originate from kinesins on protofilaments from both the 'top' and the 'bottom' of the microtubule. Due to our measurements representing a projection into the (x,y)-plane, they will cause both leftward and rightward displacement independent of the true direction. As 'center' traces show clear and maximal rightward displacement, 'between' traces show smaller leftwards and rightwards displacement and 'side' traces show close-to-zero displacement, we can conclude that during the one-head-bound state, the unbound kinesin head is displaced rightwards with respect to the kinesin's coordinate system on the microtubule.

A simple geometrical calculation concludes that the lateral microtubule position *r* and the projected displacement *d* fulfill the relation

$$\left(\frac{r}{R_{MT}}\right)^2 + \left(\frac{d}{D_{side}}\right)^2 = 1 \tag{29}$$

with $R_{MT}$ representing the microtubule radius and $D_{side}$ denoting the 3D sidewards displacement. This equation has the form of an ellipse and was used for fitting the data in Fig. 5E.

### 3 Supplementary Figures

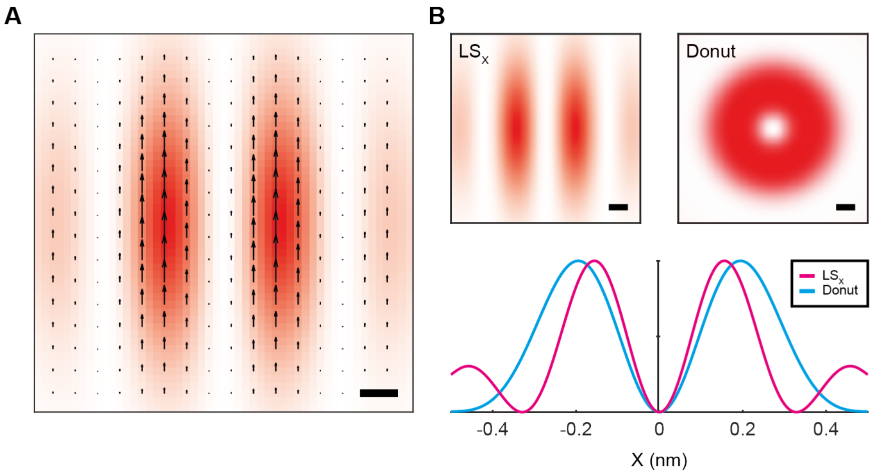


**Fig. S1. MINFLUX excitation intensity distribution and polarization vector. (A)** Profile of the excitation intensity distribution $LS_x$ in the focal plane with the corresponding lateral polarization vector illustrated by black arrows. **(B)** Comparison of the $LS_x$ and donut excitation intensity distribution in the focal plane (top) and via their normalized lateral profiles along the x-axis (bottom) showing the increased curvature of the 1D-minimum. Scale bars, 100 nm.

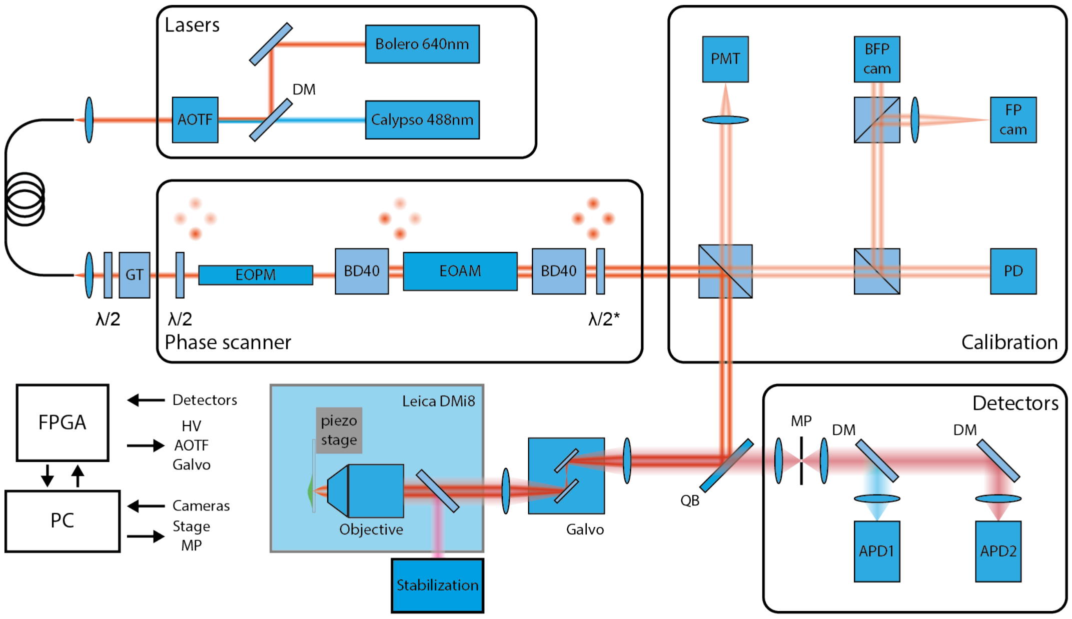


**Fig. S2. Compact arrangement of the MINFLUX setup.** Light from either a 640 nm, or 488 nm single frequency laser is split into four beamlets and phase-modulated by the phase scanner module before being guided towards the microscope body by a 90:10 beam splitter. Coarse scanning is performed by four galvanometrically steered mirrors in a quadscanner arrangement (Galvo) before focusing the beamlets into the sample. Fluorescence signal is separated from the excitation by a customized quad band filter (QB) focused onto a motorized pinhole (MP) and detected by two spectrally separated avalanche photodiodes (APDs). Calibration of the phase scanner is performed in the calibration module described in fig. S3. A stabilization unit actively locks the sample position with sub-nanometer precision as illustrated in fig. S13. (DM – dichroic mirror, AOTF – acousto optical tunable filter, GT – Glan-Thompson polarizer, EOPM – electro optical phase modulator, BD40 – polarization dependent beam displacer, $\lambda/2^*$ - segmentented half-wave plate, PMT – photomultiplier tube, BFP cam – camera in a plane equivalent to the back focal plane, FP cam – camera in a plane equivalent to the focal plane, PD – photo diode, FPGA – field-programmable gate array, HV – high voltage)

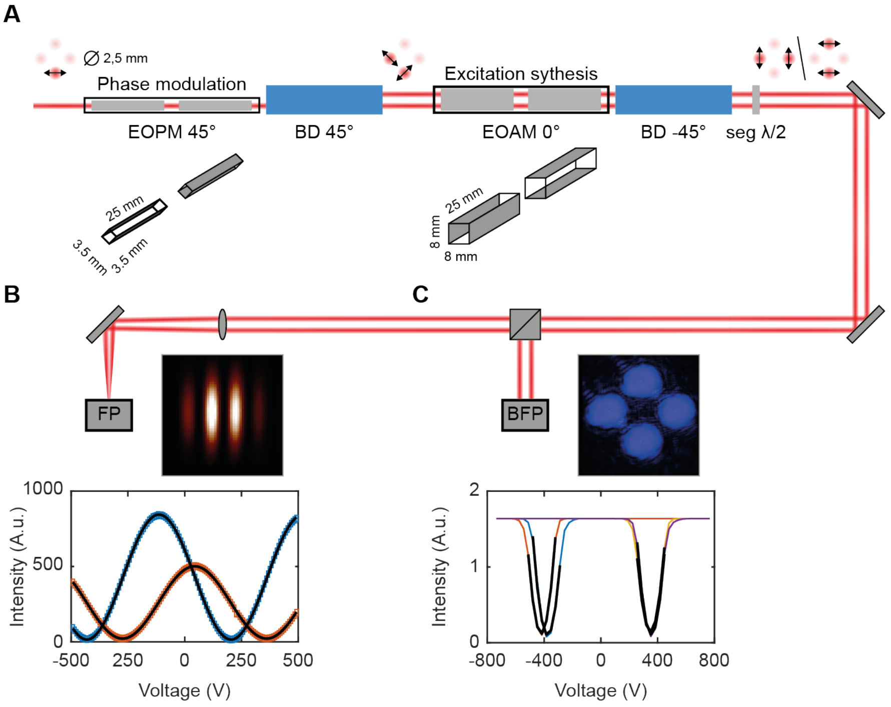


**Fig. S3. MINFLUX phase-scanner for synthesizing the excitation intensity distribution. (A)** A single horizontally polarized laser beam passes two orthogonal electro-optical crystals aligned at a 45° angle (EOPM). These crystals imposes a phase difference $\Delta\phi$ between the two polarization components along the crystal axes depending on the applied voltage which are the separated by a polarizing beam displacer (BD) also aligned at a 45° angle. The resulting two beams pass another pair of electro-optical crystals (EOAM), aligned at a 0° angle either keeping the incoming polarization state or rotating it by 90°. Afterwards, a second polarizing BD oriented at 90° to the first one generates either two horizontally separated or two vertically separated beams. Since opposite beams are orthogonally polarized a segmented $\lambda/2$-plate is installed to align these polarizations. To calibrate these electro-optical modulators, two cameras located in planes conjugated to the focal plane (FP) (B) and conjugated to the back focal plane (BFP) (C) monitor the laser beams. **(B)** The camera monitoring the focal plane is used for the calibration of the phase modulation by recording the voltage dependent intensity at the center of the focus and evaluating the frequency and phase of the recorded sine-function. **(C)** The camera monitoring the back focal plane is used for the calibration of the excitation intensity distribution synthesis by recording the voltage-dependent intensity at the position of the four beams and by evaluating the global minima.

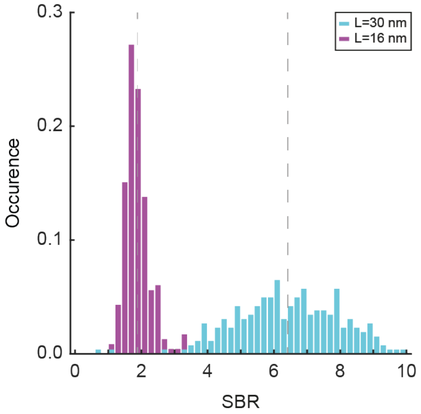


**Fig. S4. Signal-to-background ratio of surface immobilized ATTO 647N.** Normalized histogram of the measured signal-to-background ratio of surface immobilized ATTO 647N molecules using an *L* of 30 nm (turquoise) and 16 nm (purple).

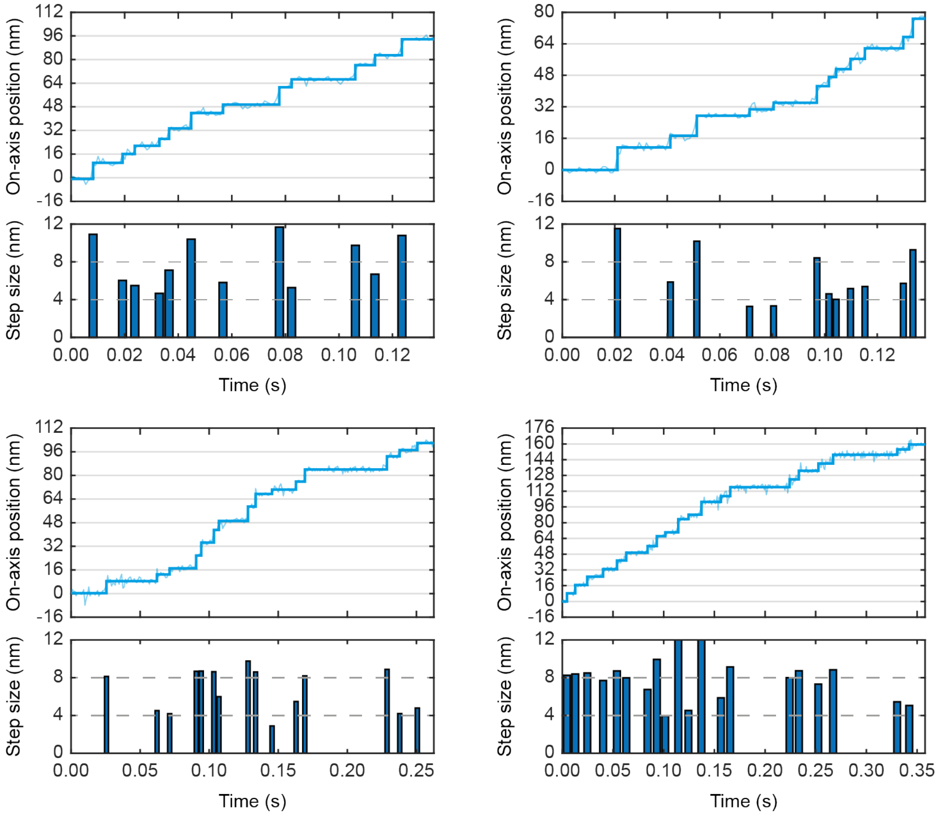


**Fig. S5. Kinesin center-of-mass substeps at physiological ATP concentration.** Exemplary on-axis position traces for the coiled-coil DOL1 labeled kinesin construct N356C recorded at 1 mM ATP concentration with the detected step function shown as thick dark lines. For each trace the corresponding step sizes are shown below with dashed grey lines marking 4 nm and 8 nm.

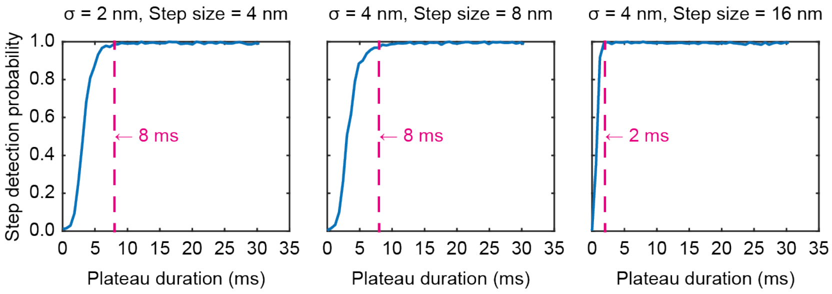


**Fig. S6. Step detection probability for various kinesin step sizes and noise levels.** Numeric simulations of the step detection probability as a function of the plateau duration for different kinesin step sizes and position noise level $\sigma$. Dashed magenta lines highlight the plateau duration at which more than 90 % of all steps are detected the step-finding algorithm.

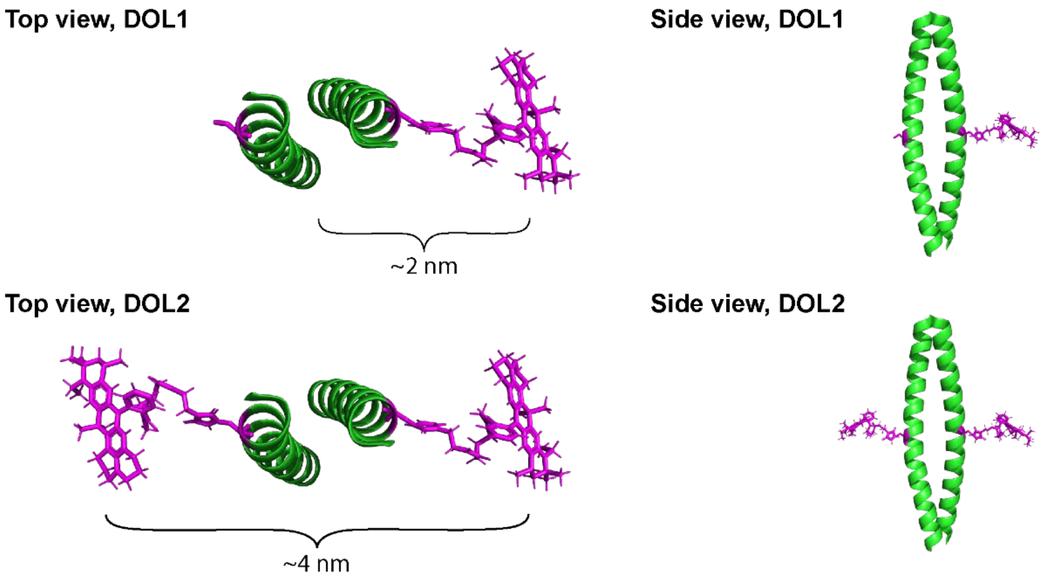


**Fig. S7. DOL1- and DOL2-labeling of the coiled-coil domain of kinesin**. Schematic illustration of the coiled-coil domain (green) labeling approach installing either one fluorophore (DOL1, yellow, top) or two fluorophores (DOL2, yellow and magenta, bottom) at amino acid position N356C. The protein structure used is from the representative parallel coiled-coil dimerization region of cortexillin I (PDB: 1D7M). The structure of ATTO 647N maleimide was inferred from its reported molecular weight, converted into a 3D structure by MM2 energy minimization in Chem3D Pro (Perkin Elmer) and added to the coiled-coil domain at XXX comparable to N356 of kinesin. The distance between the two fluorophore centers measures ~4 nm, implying a fluorophore displacement from the coiled-coil axis of ~2 nm.

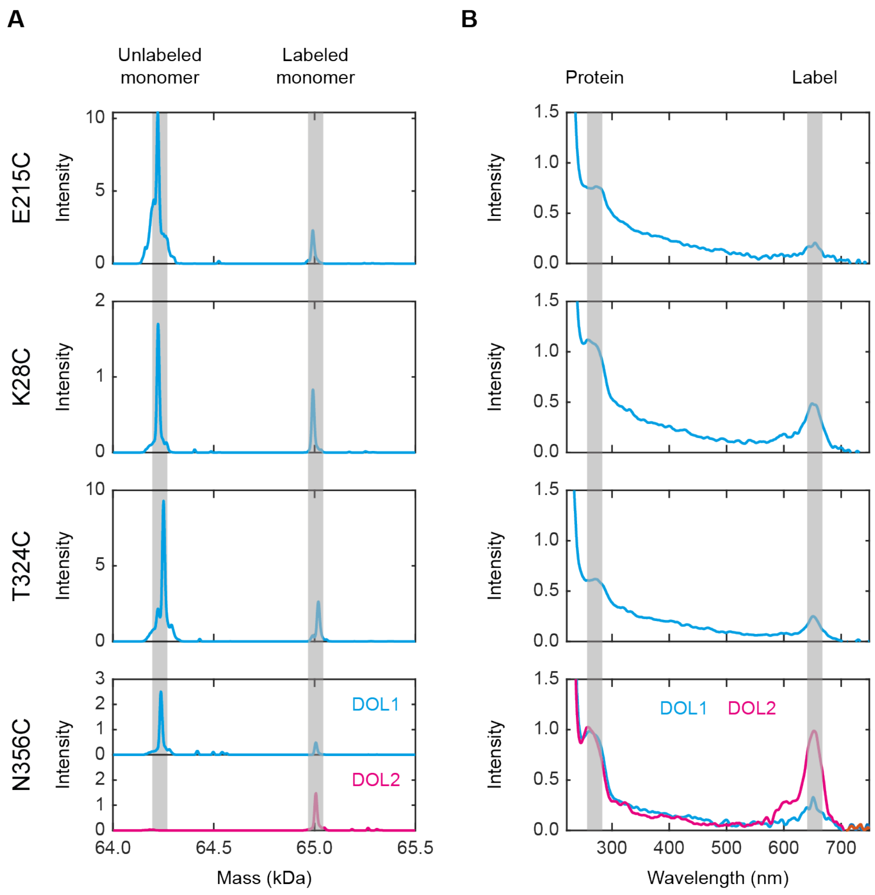


**Fig. S8. Mass and UV-Vis spectra of labeled kinesin constructs. (A)** Mass spectra showing the peaks for the labeled and unlabeled monomers of the four kinesin constructs used. **(B)** Absorption spectra in the ultra-violet (UV) and visible (Vis) spectral range. Protein absorption corresponds to the peak around 280 nm, whereas the peak around 650 nm corresponds to ATTO 647N.

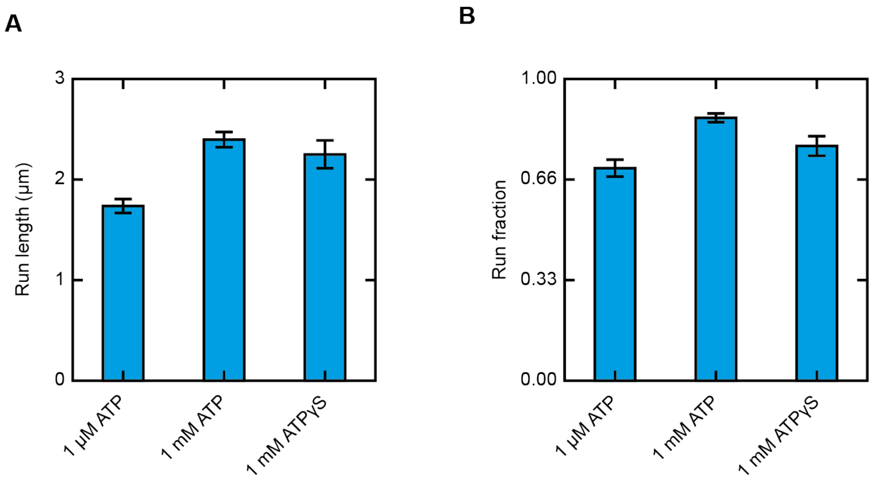


**Fig. S9. Run length and run fraction of kinesin construct T324C under different ATP conditions. (A)** Run length of kinesin construct T324C labeled with ATTO 647N maleimide as calculated from TIRF measurements at 1 µM ATP ($N_{ATP,1\mu M}$=71), 1 mM ATP ($N_{ATP,1mM}$=141) and 1 mM ATPγS ($N_{ATP\gamma S,1mM}$=57) using kymograph analysis. **(B)** Run fractions were calculated by dividing the run length by the distance between starting point and plus end of the microtubule. Bar plots display the mean and standard error of the mean.

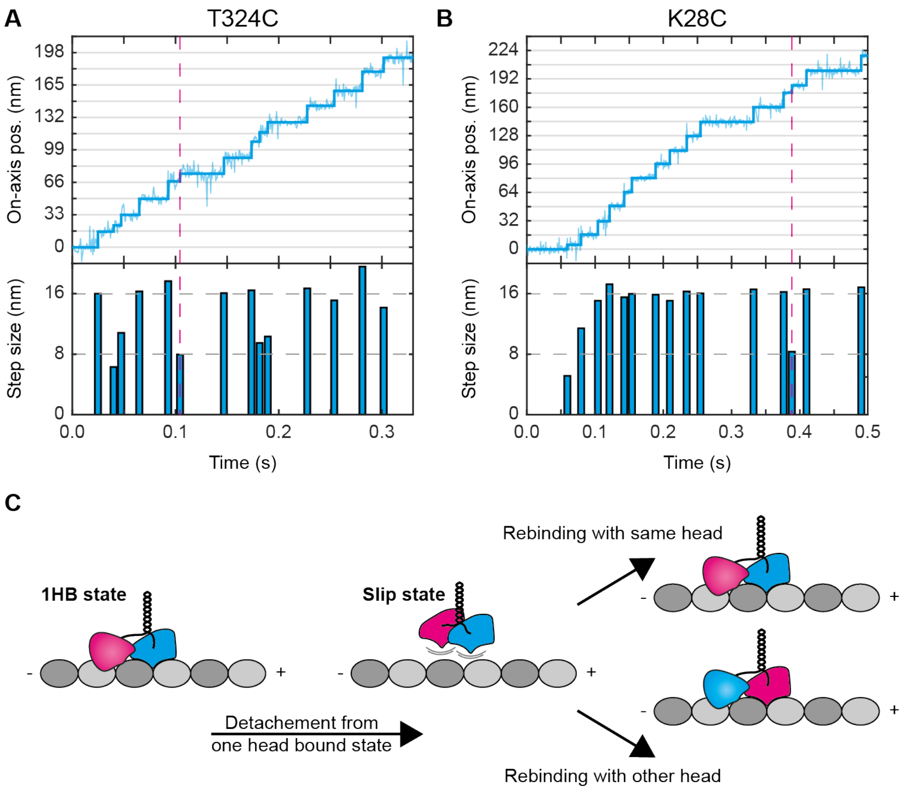


**Fig. S10. Observation of unpaired kinesin substeps explained by slip states.** Examples of an unpaired substep occurring **(A)** at around 0.1 s -within a trace recorded with the kinesin construct T324C- and **(B)** at around 0.4 s within a trace recorded with the kinesin construct K28C. Unpaired substeps are highlighted by magenta dashed lines. **(C)** Schematic illustration of kinesin detaching from the microtubule in its 1HB (cyan head bound, magenta head unbound) state and entering the suggested slip state. If kinesin rebinds to the microtubule with the same head (cyan), the expected walking pattern is recorded. If kinesin rebinds to the microtubule with its other head (magenta), unpaired substeps are detected.

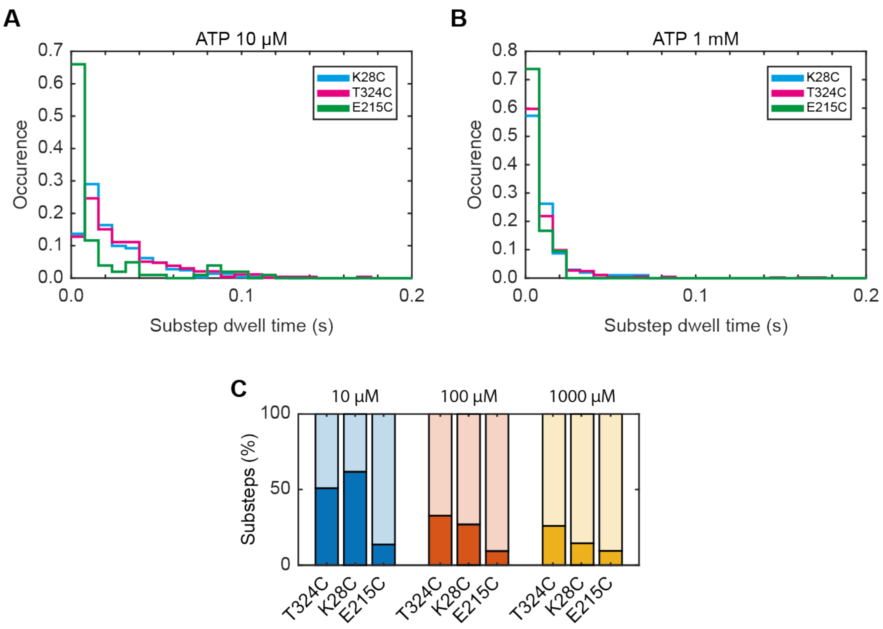


**Fig. S11. Comparison of the substep characteristics for different kinesin constructs.** Histograms of the substep dwell times recorded at **(A)** 10 μM ATP and **(B)** 1 mM ATP for the kinesin constructs K28C (cyan), T324C (magenta) and E215C (green). **(C)** Fraction of detected substeps for the kinesin constructs K28C, T324C and E215C at 10 μM (blue), 100 μM (red) and 1mM ATP (yellow) concentration.

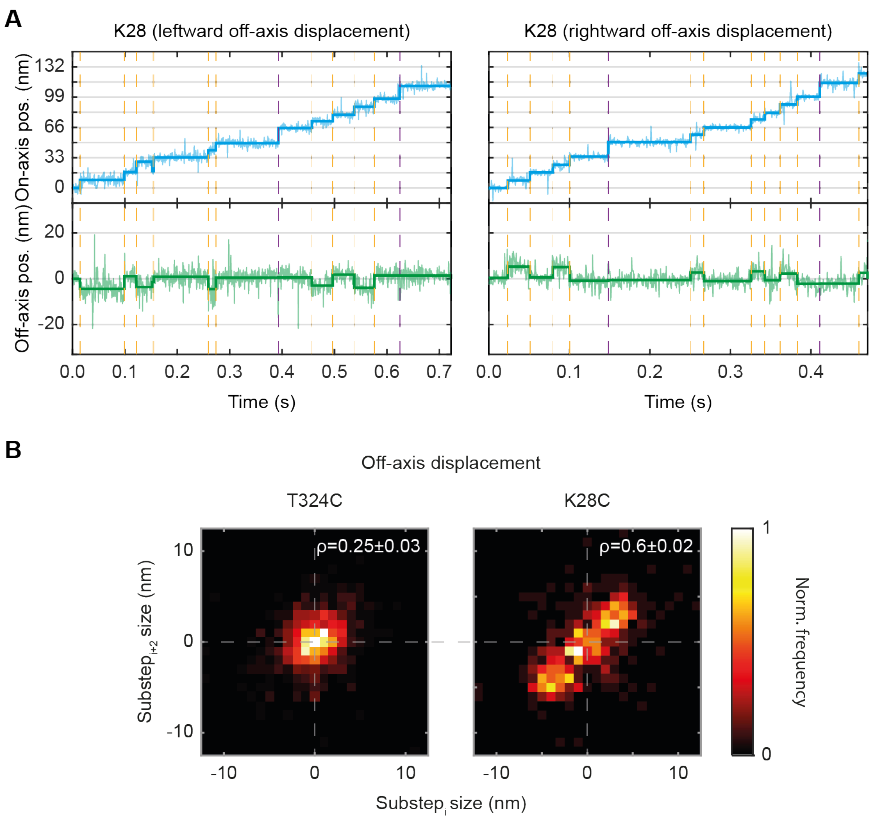


**Fig. S12. Observation of off-axis displacement during kinesin's 1HB state. (A)** Exemplary traces (top) of kinesin construct K28C recorded at 10 µM ATP showing exclusively either leftward (left) or rightward (right) off-axis displacement (bottom). The substeps that mark transitions into or out of the 1HB state are highlighted with orange dashed lines. Regular steps of 16 nm are highlighted by purple dashed lines. **(B)** 2D histogram of the off-axis displacement for successive bound to unbound and successive unbound to bound transitions showing no large displacement for kinesin construct T324C, but up to 5 nm displacement for construct K28C. For the latter, the calculated Pearson coefficient indicates that off-axis displacement occurs with similar amplitude and same sign within individual traces.

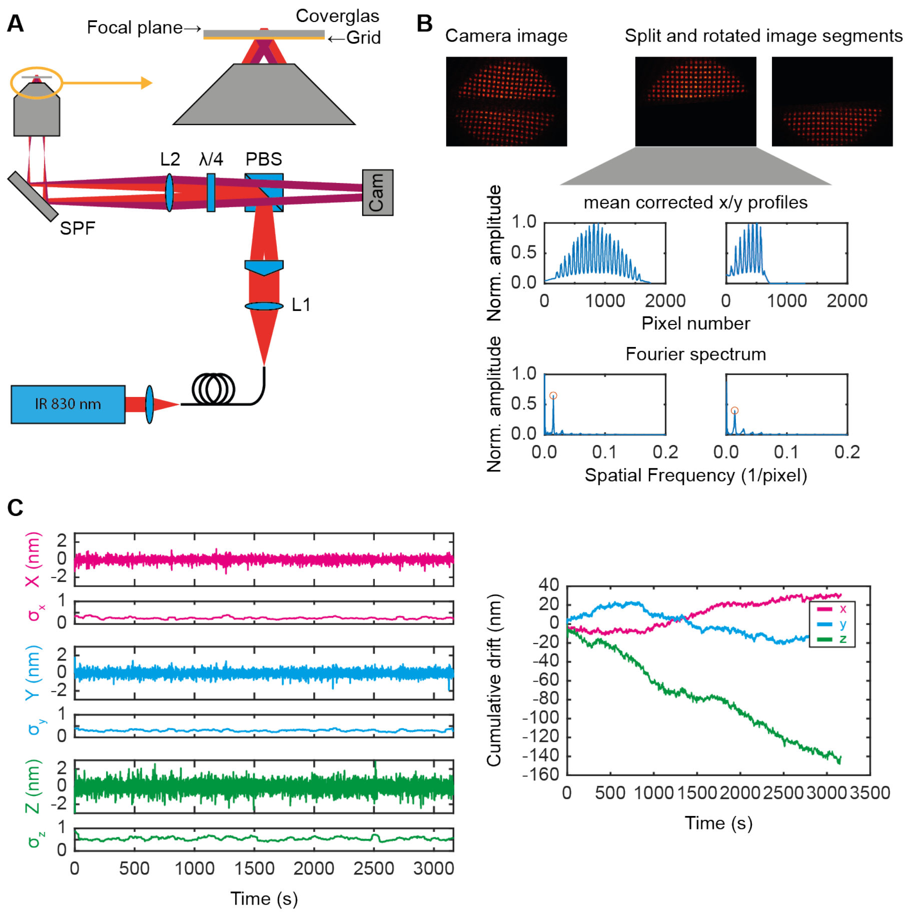


**Fig. S13. Active stabilization unit for long-term MINFLUX measurements. (A)** Simplified illustration of the optical setup of the active stabilization unit. Light from a 830 nm diode laser is fiber-coupled and expanded after the fiber before aligning the polarization horizontally with a $\lambda/2$-plate and splitting the beam into two segments by a rooftop-shaped glass plate. The two tilted beamlets are reflected by a polarizing beam splitter (PBS), sent through a $\lambda/4$-plate to achieve circular polarization and then focused (f=60 mm) in the back focal plane of the objective. The infra-red beams are coupled into the microscope body by a short pass filter (SPF). **(B)** Workflow for evaluating the 3D drift on the camera: The original camera image is split to separate the two image segments and simultaneously rotated to align the grid with the camera axes (top). For each segment the average x and y line profiles (middle) are Fourier-transformed to identify the fundamental grid frequency (bottom, highlighted in orange). The phase at this frequency is translated into the camera position of the corresponding image segment. The lateral sample drift is measured by the average drift of both image segments while drift differences between the image segments capture axial sample drift. A calibration matrix $M_{3x3}$ is used to calculate the 3D sample drift. **(C)** Left: Exemplary displacements of the sample from its reference position recorded over the course of 50 min under standard measurement conditions and the corresponding standard deviation measured by a sliding window of 1 min. Right: Cumulative 3D drift that has been corrected for throughout the measurement.

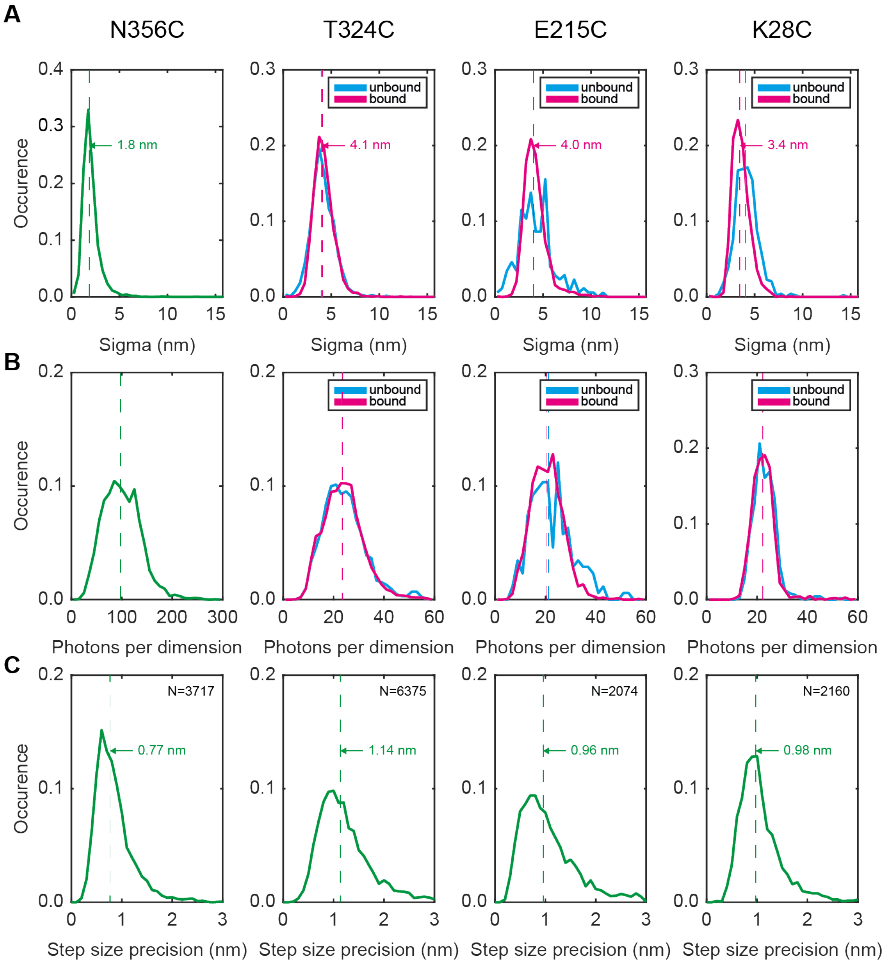


**Fig. S14. Plateau standard deviation and photon counts for all kinesin tracking measurements. (A)** Histograms of all plateau standard deviations and **(B)** histograms of all photons per localization recorded for the individual kinesin constructs. As constructs T324C, E215C and K28C allow for discrimination between the bound and unbound state of the labelled state the data plotted separately (unbound cyan, bound magenta). **(C)** Histograms of the step size precisions calculated from the plateau durations and step sizes for the individual kinesin constructs.

### 4 Supplementary Tables

**Table S1. List of all transitions included in the Hidden Markow Model with corresponding kinesin step sizes and reasoning.** B – labeled head in bound state, U – labeled head in unbound state.

| State | Stepsize | Transition | Source |
|---|---|---|---|
| 1 | 16 nm | B to B | Regular step (missed unbound state) |
| 2 | 8 nm | B to U | Substep |
| 3 | 8 nm | U to B | Substep |
| 4 | 16 nm | U to U | Missed bound state (rare) |
| 5 | 8 nm | B to B | Slip state |

**Table S2. Data assignment to the different states of the Hidden Markow Model.** Assignment was for each ATP concentration (c(ATP)) and kinesin construct, as well as for the pooled values of each kinesin construct and the pooled values of all kinesin constructs. B – labeled head in bound state, U – labeled head in unbound state.

| **Construct** | **c(ATP)** | **B→B (16 nm)** | **B→U (8 nm)** | **U→B (8 nm)** | **B→B (8 nm)** | **U→U (16 nm)** |
|---|---|---|---|---|---|---|
| T324C | 10 µM | 30% (794) | 33% (863) | 34% (883) | 1% (38) | 2% (48) |
| | 100 µM | 49% (857) | 23% (414) | 24% (429) | 2% (43) | 1% (22) |
| | 1 mM | 54% (1575) | 21% (619) | 21% (606) | 2% (63) | 1% (31) |
| | all | 44% (3226) | 26% (1896) | 26% (1918) | 2% (144) | 1% (101) |
| K28C | 10 µM | 22% (231) | 38% (396) | 38% (389) | 1% (10) | 1% (8) |
| | 100 µM | 62% (332) | 18% (97) | 18% (98) | 1% (5) | 1% (6) |
| | 1 mM | 73% (673) | 13% (116) | 13% (116) | 1% (9) | 0% (3) |
| | all | 50% (1236) | 24% (609) | 24% (603) | 1% (24) | 1% (17) |
| E215C | 10 µM | 63% (981) | 17% (265) | 17% (267) | 2% (37) | 1% (11) |
| | 100 µM | 82% (452) | 7% (39) | 7% (41) | 2% (13) | 1% (4) |
| | 1 mM | 79% (495) | 10% (61) | 10% (65) | 1% (5) | 1% (4) |
| | all | 70% (1928) | 13% (365) | 14% (373) | 2% (55) | 1% (19) |
| all | all | 51% (6390) | 23% (2870) | 23% (2894) | 2% (223) | 1% (137) |

**Table S3. MINFLUX measurement parameters.** L value, Laserpower going into the Galvo given as the mean of all powers used weighted by the number of traces recorded at each power (minimal power, maximal power), duration of a full localization $\tau$ given as the mean of all durations used weighted by the number of traces recorded at each duration (minimal duration, maximal duration)

| **Sample** | **L** | **Power mean (min, max)** | **$\tau$ mean (min, max)** |
|---|---|---|---|
| ATTO 647N-bt (2D) | 16 nm | 1.28 mW (1.17 mW, 1.37 mW) | 631 µs (631 µs, 631 µs) |
| ATTO 647N-bt (2D) | 30 nm | 0.78 mW (0.78 mW, 0.78 mW) | 631 µs (631 µs, 631 µs) |
| Kinesin N356C (1D) | 30 nm | 1.15 mW (1.04 mW, 1.63 mW) | 838 µs (616 µs, 1216 µs) |
| Kinesin T324C (2D) | 30 nm | 0.48 mW (0.23 mW, 0.59 mW) | 633 µs (607 µs, 1207 µs) |
| Kinesin E215C (2D) | 30 nm | 0.57 mW (0.52 mW, 1.04 mW) | 611 µs (607 µs, 631 µs) |
| Kinesin K28C (2D) | 30 nm | 0.59 mW (0.52 mW, 1.56 mW) | 631 µs (631 µs, 631 µs) |
